# DARPP-32 promotes ERBB3-mediated resistance to molecular targeted therapy in EGFR-mutated lung adenocarcinoma

**DOI:** 10.1101/2021.02.12.430856

**Authors:** Sk. Kayum Alam, Yongchang Zhang, Li Wang, Zhu Zhu, Christina E. Hernandez, Yuling Zhou, Nong Yang, Jian Lei, Xiaoyan Chen, Liang Zeng, Mark A. Klein, Luke H. Hoeppner

## Abstract

While molecular targeted therapies have improved prognoses of advanced stage lung adenocarcinoma expressing oncogenic driver mutations, acquired therapeutic resistance continues to be a major problem. Epidermal growth factor receptor (EGFR) activating mutations are among the most common targetable genetic alterations in lung adenocarcinoma, and EGFR tyrosine kinase inhibitors (TKIs) are recommended first-line therapy for EGFR mutation positive cancer patients. Unfortunately, most patients develop resistance to EGFR TKIs and rapid disease progression occurs. A better mechanistic understanding of therapy refractory cancer progression is necessary to develop new therapeutic approaches to predict and prevent acquired resistance to EGFR TKIs. Here, we identify a new mechanism of ERBB3-mediated resistance to EGFR TKIs in human lung adenocarcinoma. Specifically, we show that dopamine and cyclic AMP-regulated phosphoprotein, Mr 32000 (DARPP-32) physically recruits ERBB3 to EGFR to mediate a switch from EGFR homodimers to EGFR:ERBB3 heterodimers to bypass EGFR TKI-mediated inhibition to potentiate ERBB3-dependent activation of oncogenic AKT and ERK signaling that drives therapy refractory tumor cell survival. In a cohort of paired tumor specimens derived from 30 lung adenocarcinoma patients before and after the development of EGFR TKI refractory disease progression, we reveal that DARPP-32 as well as kinase-activated EGFR and ERBB3 proteins are overexpressed upon acquired EGFR TKI resistance. In vivo studies suggest that ablation of DARPP-32 protein activity sensitizes gefitinib-resistant lung tumor xenografts to EGFR TKI treatment, while DARPP-32 overexpression increases gefitinib-refractory lung cancer progression in gefitinib-sensitive lung tumors orthotopically xenografted into mice. Taken together, our findings introduce a DARPP-32-mediated, ERBB3-dependent mechanism used by lung tumor cells to evade EGFR TKI-induced cell death, potentially paving the way for the development of new therapies to prevent or overcome therapy-refractory lung adenocarcinoma progression.

## Introduction

Lung cancer is the leading cause of cancer deaths in the United States and worldwide^1^. Non-small cell lung cancer (NSCLC) represents 80-85% of lung cancer diagnoses, most of which presents as advanced disease with poor prognoses: 1-year survival rates of ~15% and median overall survival of less than 12 months^2^. Targeted therapies for NSCLC expressing oncogenic driver mutations have improved prognoses. The approximately 30% of advanced NSCLC patients with epidermal growth factor receptor (EGFR) mutations^3^ benefit from treatment with EGFR tyrosine kinase inhibitors (TKIs)^4, 5^. Unfortunately, most patients develop resistance to EGFR TKIs and rapid disease progression occurs^6, 7^. The 2019 novel coronavirus (COVID-19) pandemic has worsened the health consequences of lung cancer^8^. Lung cancer patients have a higher incidence of COVID-19 and more severe symptoms than cancer-free individuals^9-11^, which amplifies the urgency and need for improved lung cancer therapies, including new approaches to prevent or overcome therapy refractory cancer progression.

EGFR is a member of the ERBB family of receptor tyrosine kinases that bind extracellular growth factors to mediate intracellular signaling^12^. EGFR regulates cell proliferation, survival, differentiation and migration through activation of several signal transduction pathways, including the phosphatidylinositol-3 kinase (PI3K)–AKT-mammalian target of rapamycin (mTOR) cascade, mitogen-activated protein kinase (MAPK) signaling, phospholipase Cy (PLCγ)–protein kinase C (PKC) pathway, and Janus kinase 2 (JAK2)–signal transducer and activator of transcription 3 (STAT3) signaling^13, 14^. EGFR was identified as promising therapeutic molecular target based on its overexpression and correlation with poor prognosis in NSCLC^15^. Consequently, two small molecule inhibitors targeting EGFR, gefitinib (Iressa®, 2003) and erlotinib (Tarceva®, 2004), received FDA approval as treatment for NSCLC patients who had failed chemotherapy^15^. 10% of NSCLC patients treated with EGFR inhibitor responded, mostly women, non-smokers, East Asians and patients with adenocarcinomas displaying bronchioloalveolar histology. Molecular studies revealed responders typically possessed EGFR mutations. In-frame deletions of amino acids 747-750 in exon 19 made up 45% of the EGFR mutations and 40-45% consisted of L858R mutations in exon 21 of EGFR^4, 5^. These activating mutations hyperactivate the kinase activity of EGFR to stimulate oncogenic signaling that promotes tumor cell survival, proliferation, differentiation and migration^16-18^. EGFR tyrosine kinase inhibitors (TKIs) are the recommended first-line therapy for NSCLC patients positive for an EGFR mutation based on trials with gefitinib, erlotinib, and afatinib showing significant improvements in response rate and progression free survival compared with first-line chemotherapy^19^. Although EGFR mutation positive patients respond well to first-line EGFR TKIs, NSCLC inevitably progresses in most patients after 9-12 months^20^. In 40-60% of these patients, an exon 20 T790M mutation occurs in EGFR^21, 22^. Osimertinib, a third-generation EGFR TKI targeting the T790M mutation and the primary activating EGFR mutations, can be used to overcome resistance to the first-generation TKIs and has also been recently approved in the United States as a first-line treatment for advanced NSCLC patients with EGFR sensitizing mutations^23, 24^. However, only ~60% of patients with T790M mutations respond to osimertinib, and in those responding patients, NSCLC progression typically occurs in less than 10 months^23^. Osimertinib resistance mechanisms include EGFR C797S mutations and histological/phenotypic transformation^25, 26^. Furthermore, acquired resistance to EGFR TKIs can develop through activation of other oncogenic pathways, such as c-Met amplification, activation of the PI3K/AKT pathway, and EGFR-independent phosphorylation of ERBB^27^. Thus, better treatment options to overcome EGFR TKI resistance are necessary.

We recently have identified the role of dopamine signaling in lung cancer^28-30^. In particular, we have shown that cyclic AMP-regulated phosphoprotein, Mr 32000 (DARPP-32), and its N-terminal truncated isoform named t-DARPP, contribute to lung oncogenesis^28, 30^. Here, we demonstrate DARPP-32 and t-DARPP proteins promote EGFR TKI refractory disease progression. DARPP-32, and its transcriptional splice variant t-DARPP, are frequently overexpressed in breast, gastric, thoracic, colon, pancreatic, and other adenocarcinomas, where their aberrant upregulation contributes to oncogenesis through regulation of cellular processes, including proliferation, survival, migration, and angiogenesis^30-38^. DARPP-32 was initially discovered as an effector of dopaminergic neurotransmission and as a substrate of dopamine-activated protein kinase A (PKA)^39^. Phosphorylation at T34 by PKA causes DARPP-32-mediated inhibition of protein phosphatase-1 (PP-1)^40^. DARPP-32 is converted to an inhibitor of PKA upon phosphorylation of its T75 residue by cyclin-dependent kinase 5 (Cdk5)^41^. The ability of DARPP-32 to function as either a kinase or a phosphatase inhibitor enables it to precisely modulate dopaminergic neurotransmission^41, 42^. In the early 2000s, El-Rifai and colleagues discovered that DARPP-32 is frequently amplified and upregulated in gastric cancer^32, 35^. Cloning and sequence assembly revealed a novel transcriptional splice variant of DARPP-32 is also overexpressed in gastric cancer. The N-terminally truncated isoform of DARPP-32, termed t-DARPP, was found to utilize a unique alternative first exon located within intron 1 of *phosphoprotein phosphatase-1 regulatory subunit 1B (PPP1R1B),* the gene that transcribes DARPP-32 and t-DARPP proteins^35^. t-DARPP lacks the first 36 amino acids of DARPP-32, including the T34 phosphorylation residue required for DARPP-32-mediated PP-1 inhibition^35^. Elevated expression of t-DARPP isoform in NSCLC is associated with poor overall survival and increasing tumor (T) stage^30^. Our findings presented in this report suggest that overexpression of DARPP-32 isoforms in EGFR-mutated NSCLC promotes EGFR:ERBB3 “bypass signaling” that enables tumor cells to evade EGFR TKI monotherapy-induced apoptosis by potentiating oncogenic AKT and ERK signaling.

## Methods

### Cell culture

Gefitinib-sensitive HCC827 parental (HCC827P) and HCC827 gefitinib-resistant (HCC827GR) human NSCLC cell lines were a generous gift from Dr. Pasi A. Jänne at Dana-Farber Cancer Institute^43^. Gefitinib-sensitive PC9 parental (PC9P) and its gefitinib-resistant counterparts (PC9GR2, PC9GR3) were kindly provided by Dr. Aaron N. Hata at Massachusetts General Hospital^44^. HCC827P and PC9P cells were grown in RPMI-1640 medium (Corning). RPMI-1640 medium containing 1 μm of gefitinib (Selleckchem) was used to culture HCC827GR, PC9GR2, and PC9GR3 cells. HEK-293T cells were purchased from American Type Culture Collection (ATCC) and maintained in Dulbecco’s modified Eagle’s medium (DMEM; Corning). Culture medium for each of the cell lines was supplemented with 10% fetal bovine serum (FBS; Millipore), 1% Penicillin/Streptomycin antibiotics (Corning), and 25 μg/mL Plasmocin prophylactic (Invivogen). All cell lines were authenticated by their source and were subsequently routinely authenticated via morphologic inspection.

### Generation of stable cell lines

Human DARPP-32 and t-DARPP cDNAs cloned in pcDNA3.1 were kindly provided by Dr. Wael El-Rifai at University of Miami Health System^33^. To generate retrovirus, we first subcloned FLAG-tagged DARPP-32 and t-DARPP cDNAs into pMMP vectors that were a kind gift from Dr. Debabrata Mukhopadhyay at Mayo Clinic in Jacksonville, Florida^45^. HEK-293T cells were next transfected with pMMP vectors together with retrovirus packaging plasmids using Effectene transfection reagents (Qiagen) according to the manufacturer’s protocol. Two days after transfection, the retrovirus was collected from the cell culture medium, concentrated using Retro-X concentrator (Takara), and used immediately to transduce human NSCLC cell lines, HCC827P and PC9P, as described previously^28^.

To prepare lentivirus, LacZ shRNA (control) and 4 different DARPP-32 shRNAs cloned in pLKO.1 plasmids (Sigma) along with their corresponding packaging plasmids were transfected in human HEK-293T cells. The lentivirus was isolated from cell culture medium 48h after transfection, concentrated using Lenti-X concentrator (Takara), and used immediately to transduce HCC827GR, PC9GR2, and PC9GR3 lung cancer cell lines, as reported previously^28^. Stable DARPP-32 knockdown cells were used for experiments following 72h of puromycin (Sigma) selection.

HCC827P and HCC827GR cells transduced with lentivirus containing the luciferase gene were used to determine tumor growth in orthotopic murine models. Briefly, luciferase gene encoding lentivirus was prepared by transfecting MSCV Luciferase PGK-hygro plasmids obtained through Dr. Scott Lowe via Addgene (#18782) along with their corresponding packaging plasmids into human HEK-293T cells. Two days post-transfection, the lentivirus collected from the cell culture media was concentrated using Retro-X concentrator (Takara). The concentrated lentivirus was used immediately to transduce human NSCLC cell lines, HCC827P and HCC827GR, as described previously^46^. Luciferase-labeled stable human NSCLC cells were obtained following 72h of hygromycin (Sigma) selection after transduction.

### Antibodies

For detection of proteins in immunoblotting experiments, we purchased monoclonal antibodies (1 μg/μl) against phosphorylated EGFR (Y1068; Cat no.: 3777; Dilution 1:1000), total EGFR (Cat no.: 4267, Dilution 1:1000), phosphorylated ERBB2 (Y1221/1222; Cat no.: 2243; Dilution 1:1000), total ERBB2 (Cat no.: 4290, Dilution 1:1000), phosphorylated ERBB3 (Y1289; Cat no.: 2842; Dilution 1:1000), total ERBB3 (Cat no.: 12708, Dilution 1:1000), PARP-I (Cat no.: 9542; Dilution 1:1000), Caspase-3 (Cat no.: 9662; Dilution 1:1000), Cleaved Caspase-3 (Cat no.: 9664; Dilution 1:1000), phosphorylated AKT (S473; Cat no.: 4060; Dilution 1:1000), total AKT (Cat no.: 4691; Dilution 1:1000), phosphorylated ERK1/2 (T202/Y204; Cat no.: 4370; Dilution 1:1000), total ERK1/2 (Cat no.: 4695; Dilution 1:1000) from Cell Signaling Technology. Monoclonal antibodies (200 μg/ml) to detect DARPP-32 (Cat no.: sc-135877; Dilution 1:200) and α-Tubulin (Cat no.: sc-5286; Dilution 1:1000) protein were obtained from Santa Cruz Biotechnology. Horseradish peroxidase (HRP)-conjugated secondary antibodies (1 μg/μl) purchased from Cell Signaling Technology were used to detect primary antibodies raised in either rabbit (Cat no.: 7074; Dilution 1:5000) or mouse (Cat no.: 7076; Dilution 1:5000).

### Immunoblotting

Radioimmunoprecipitation assay (RIPA) buffers (Millipore) containing protease (Roche) and phosphatase inhibitors (Millipore) were used to lyse human NSCLC cell lines. Proteins quantified using the Quick Start Bradford protein assay reagents (Bio-Rad) were separated via 4-20% gradient SDS-PAGE (Bio-Rad) and transferred to polyvinyl difluoride membranes (PVDF; Millipore). Membranes blocked with 5% bovine serum albumin (BSA; Sigma-Aldrich) were then incubated with primary and corresponding secondary antibodies overnight and for 2h, respectively. Enzyme-based chemiluminescence substrate (Thermo Fisher Scientific) was used to detect antibody-reactive protein bands.

### Cell survival assay

Human NSCLC cell lines, HCC827P and HCC827GR, each plated in a 96-well microplate at a concentration of 5000 cells/well, were used to determine cell survival in the presence of increasing concentrations of gefitinib. Seventy-two hours post-gefitinib treatment, cell viability was assessed using MTS1-based CellTiter 96® AQueous One System (Promega). Absorbance of cell culture medium recorded at 490 nm using a Synergy Neo2 microplate spectrophotometer (Biotek) was used to calculate IC_50_ values of gefitinib in different experimental groups.

### Apoptosis analysis

To determine gefitinib-induced cell apoptosis, 1x10^5^ human EGFR-mutated NSCLC cells plated in 60-mm dishes were stained with fluorescein isothiocyanate (FITC)-conjugated anti-Annexin V antibodies (BD Biosciences) together with propidium iodide (BD Biosciences) following 24h gefitinib treatment. The number of early apoptotic cells (Annexin-positive and propidium-iodide-negative) determined by flow cytometry-based analysis was counted to measure apoptotic cell death.

### Immunofluorescence

PC9P and PC9GR3 cells fixed in 4% paraformaldehyde (Boston Bioproducts) were permeabilized in cold methanol (Fisher). Permeabilized cells were then used for immunofluorescence staining using primary antibodies against phosphorylated-EGFR (Y845; BD Biosciences; Cat No.: 558381; Dilution 1:200) and phosphorylated-ERBB3 (Y1289; Cell Signaling Technology; Cat No.: 2842; Dilution 1:200). Secondary antibody staining was performed by incubation with matching Alexa Fluor 488-conjugated anti-rabbit antibody (Molecular Probes; Cat no.: A11008; Dilution 1:400) and Alexa Fluor 594-conjugated anti-mouse antibody (Molecular Probes; Cat no.: A11005; Dilution 1:400). Cell nuclei were stained using 4’, 6-diamidino-2-phenylindole, dihydrochloride (DAPI), including ProLong® Gold Antifade Reagent (Cell Signaling Technology). Images captured in Zeiss Apotome.2 microscope (63X objective, 1.25 NA) were processed using ZEN microscope software (Zeiss). The average fluorescence intensity for green and red signals was calculated using ZEN microscope software and reported.

### Immunoprecipitation

RIPA buffers (Millipore) supplemented with protease (Roche) and phosphatase inhibitors (Millipore) were used to lyse human NSCLC cell lines for immunoprecipitation studies. Bradford-based protein assay (Bio-Rad) was used to determine the concentration of harvested cell lysates and 500 μg of protein lysate was loaded into the supplied spin column (Catch and Release Immunoprecipitation Kit; Millipore). Immunoprecipitation using antibodies against FLAG (Cell Signaling Technology; Cat No.: 14793), ERBB3 (Cell Signaling Technology; Cat No.: 12708), and DARPP-32 (Santa Cruz Biotechnology; Cat No.: sc-135877) was achieved by following manufacturer’s protocol (Cat no.:17-500; Millipore).

### Proximity ligation assay

Human lung adenocarcinoma PC9P and PC9GR3 cells seeded in chamber slides at a density of 1x10^4^ cells/well were incubated with or without 100nM gefitinib for 24h. Cells were then washed with PBS (Corning), fixed with 4% paraformaldehyde (Boston Bioproducts), and permeabilized in cold methanol (Fisher). Permeabilized cells were incubated with primary antibodies against phosphorylated-EGFR (Y845; BD Biosciences; Cat No.: 558381; Dilution 1:200) and phosphorylated-ERBB3 (Y1289; Cell Signaling Technology; Cat No.: 2842; Dilution 1:200) diluted in SignalStain® Antibody Diluent (Cell Signaling Technology). Proximity ligation assay (PLA) probes designed to bind to corresponding primary antibodies were ligated, amplified, and washed according to the manufacturer’s instructions. Untreated lung cancer cells incubated with rabbit (Cell Signaling Technology, Cat No.: 2729) and mouse non-immune IgG (Cell Signaling Technology, Cat No.: 5415) were used for negative controls. Slides mounted in ProLong® Gold Antifade Reagent with DAPI (Cell Signaling Technology) were imaged using a Zeiss Apotome.2 microscope (20X objective, 0.8 NA) and processed using ZEN microscope software (Zeiss). PLA results were quantified by counting the number of red PLA signals normalized to the total number of DAPI-stained nucleus using Image J software (Version 1.6.0_24; https://imagej.nih.gov/ij). The average number of PLA signals per cells in 6-10 random microscopic fields for each sample was recorded.

### Gene expression analyses

Information regarding EGFR mutations and *PPP1R1B* gene expression in NSCLC were obtained from The Cancer Genome Atlas database (TCGA), an open access database that is publicly available at http://www.cbioportal.org^47, 48^. We selected TCGA Pan Cancer cohort^49^ as our data source, which contains detailed information about DNA mutations, copy-number changes, mRNA expression, gene fusions, and DNA methylation of 9,125 tumors. Mutation as well as mRNA expression (i.e. RNA-seq) data in 510 non-small cell lung adenocarcinoma (LUAD) patients were downloaded from cBioPortal website and classified based on the presence of EGFR mutation. We next subdivided EGFR-mutated patient cohort (n=80) between DARPP-32 low (n=76) vs high (n=4) group based on the *PPP1R1B* mRNA expression. The mRNA expression of PI3K/AKT/mTOR downstream targets (i.e. *RPS6KB1* and *RPS6KB2*) in 80 EGFR-mutated patients were obtained by merging the RNA-seq data according to the unique patient ID, such as “TCGA-05-4244”. Briefly, RNA-seq results were downloaded from cBioPortal website by entering gene symbols. Downloaded results were then sorted based on the patient ID. Normalized mRNA expression of *RPS6KB1* and *RPS6KB2* (i.e. Z-score) were reported in the box plot graph.

Proteome and phosphoproteome data generated from RPPA study in NSCLC patients (n=360) were extracted from the TCGA Pan Cancer study. Based on the information about EGFR mutation and *PPP1R1B* mRNA expression, we divided EGFR mutant patient cohort (n=41) between DARPP-32 low (Z-score<1.5; n=36) vs high (Z-score≥1.5; n=5) group. Similar to previous analysis, normalized protein and phosphoprotein expressions of PI3K/AKT/mTOR were calculated and reported in the box plot graph.

### *In vivo* orthotopic lung cancer model

Six to eight-week-old pathogen-free SCID/NCr mice purchased from the Charles River Laboratories were maintained in accordance with protocols approved by the University of Minnesota Institutional Animal Care and Use Committee (IACUC). Mice were allowed one week to acclimate to their surroundings, bred, maintained under specific pathogen-free (SPF) conditions in a temperature-controlled room with alternating 12h light/dark cycles, and fed a standard diet. Eight- to twelve-week-old male and female mice were anesthetized with pharmaceutical grade ketamine (90-120 mg/kg) and xylazine (5-10 mg/kg) via intraperitoneal injection under a laminar flow hood in an SPF room within the animal facility. Each fully anesthetized mouse was placed in the right lateral decubitus position and the left lateral chest was sterilized. One-million luciferase-labeled human HCC827GR and HCC827P lung cancer cells suspended in 80 μl PBS and high concentration Matrigel (Corning; Cat. no.: 354248) were orthotopically injected in the left thoracic cavity of each mouse. Based on the captured luminescence images of mice using an In-Vivo Xtreme xenogen imaging system (Bruker) as described^28^, mice were randomly divided into two groups with nearly same average luminescence intensity. After establishment of the lung tumor, mice were administered either vehicle or gefitinib (25mg/Kg) every other day for 2 weeks. Upon completion of the study, mice were euthanized using asphyxiation by CO2 inhalation to effect with a flow rate displacing less than 30% of the chamber volume per minute in accordance with IACUC euthanasia guidelines and consistent with recommendations of the Panel of Euthanasia of the American Veterinary Medical Association. Following euthanasia, lungs were perfused, harvested, and portions the lungs from each mouse were preserved in formalin for immunohistochemistry, *RNAlater* Stabilization Solution (Thermo Fisher Scientific) for RNA extraction, and flash frozen for protein extraction. Bruker molecular imaging software was used to calculate luciferase intensity (total photons count) of tumor cells in each mouse. Tumour growth was determined by plotting average luciferase intensity over time in GraphPad Prism 8 software.

### *In vivo* subcutaneous lung tumor model

Eight- to twelve-week-old male and female mice were subcutaneously injected with 2×10^6^ luciferase-labeled human PC9P lung cancer cells suspended in 80 μl PBS and high concentration Matrigel. To determine tumor growth, the tumor volume was measured every week using the formula: (length x width^2^)/2. After establishment of palpable tumor (≥150mm^3^), mice were randomly divided into two groups and administered either vehicle or gefitinib (25mg/Kg). At the endpoint, mice were euthanized by CO2 asphyxiation. Extirpated tumors were photographed, weighed, and preserved in formalin for immunohistochemistry analysis. This study was performed in accordance with approved University of Minnesota IACUC protocols.

### Clinical workflow and patient selection

Patients who met the following criteria were enrolled in this study: (1) pathologically confirmed advanced lung adenocarcinoma; (2) NGS identified EGFR exon 21 L858R mutation; (3) treatment with gefinitib or erlotinib in the first-line setting; and (4) accessed with disease progression and available tumor sample at baseline and progression. Patients were examined every two weeks after EGFR TKI administration and 20% incensement of tumor burden is considered as disease progression according to RECIST 1.1. Pathological diagnosis and staging were carried out according to the staging system of the 2009 International Association for the Study of Lung Cancer (version 8). Written informed consent was obtained from all the patients prior to inclusion to this study. Approval was also obtained from Hunan Cancer Hospital Institutional Review Board (IRB) Committee.

### Immunohistochemistry

Immunohistochemistry using primary antibodies against Ki-67 (Cell Signaling Technology, Cat. No.: 12202, Dilution 1:100), DARPP-32 (Abcam, Cat. No.: ab40801, Dilution 1:100), EGFR (Abcam, Cat. No.: ab52894, Dilution 1:50), p-EGFR (Abcam, Cat. No.: ab40815, Dilution 1:100), ERBB3 (Abcam, Cat. No.: ab85731, Dilution 1:50), p-ERBB3 (ab101407, Dilution 1:200) was performed on formalin-fixed, paraffin-embedded lung tumor tissue specimens derived from mice (for Ki-67) or lung adenocarcinoma patients (for the other antibodies). For evaluation of the morphology, 5 μm sections were stained with hematoxylin and eosin. After dewaxing, hydration, and antigen retrieval, primary antibody-treated slides were incubated with secondary antibody (Cell Signaling Technology). Finally, the degree of staining of the tissue specimens was observed under the Zeiss Apotome.2 microscope (20X objective, 0.8 NA) after DAB staining (Cell Signaling Technology) and hematoxylin (IHC World) counterstaining.

### Statistics

To compare differences between two groups, two-way unpaired t-test was performed and values of *P* ≤0.05 were considered significant. One-way analysis of variance (ANOVA) followed by Dunnett’s test was used to determine statistically significant differences between multiple groups (greater than two). Data expressed as mean ±SEM are representative of at least three independent experiments.

### Data availability

The authors declare that the data supporting the findings of this study are available within the article and its supplementary information.

## Results

### DARPP-32 is upregulated in EGFR TKI-resistant NSCLC cells

Given the ability of DARPP-32 to modulate oncogenic signaling^50, 51^, we hypothesized that DARPP-32 contributes to acquired EGFR TKI resistance in NSCLC. To test this hypothesis, we utilized two well-characterized NSCLC models of EGFR TKI resistance. Gefitinib-resistant HCC827 (EGFR^*ΔE746-A750*^) human NSCLC cells were previously generated through six months of exposure to increasing concentrations of gefitinib and shown to have acquired gefitinib resistance through a c-MET amplification^43^. Secondly, we relied on gefitinib-sensitive PC9 (EGFR^*L858R*^) human NSCLC cells and their corresponding PC9 gefitinib-resistant (PC9GR2 and PC9GR3) counterparts, which acquired gefitinib resistance through a secondary EGFR^*T790M*^ mutation following prolonged parental cell exposure to this first-generation EGFR TKI^44^. We observed reduced DARPP-32 protein levels in gefitinib-sensitive, HCC827 parental (HCC827P) and PC9 parental (PC9P) cells upon treatment with EGFR inhibitor, gefitinib (Fig. 1a-b; Supplementary Fig. 1a-b). Based on this result, we sought to examine DARPP-32 protein levels in gefitinib-resistant EGFR-mutated NSCLC cells. By immunoblotting, we observed elevated DARPP-32 protein expression in gefitinib-resistant cells relative to parental counterparts (Supplementary Fig. 2a-b).

**Figure 1:**
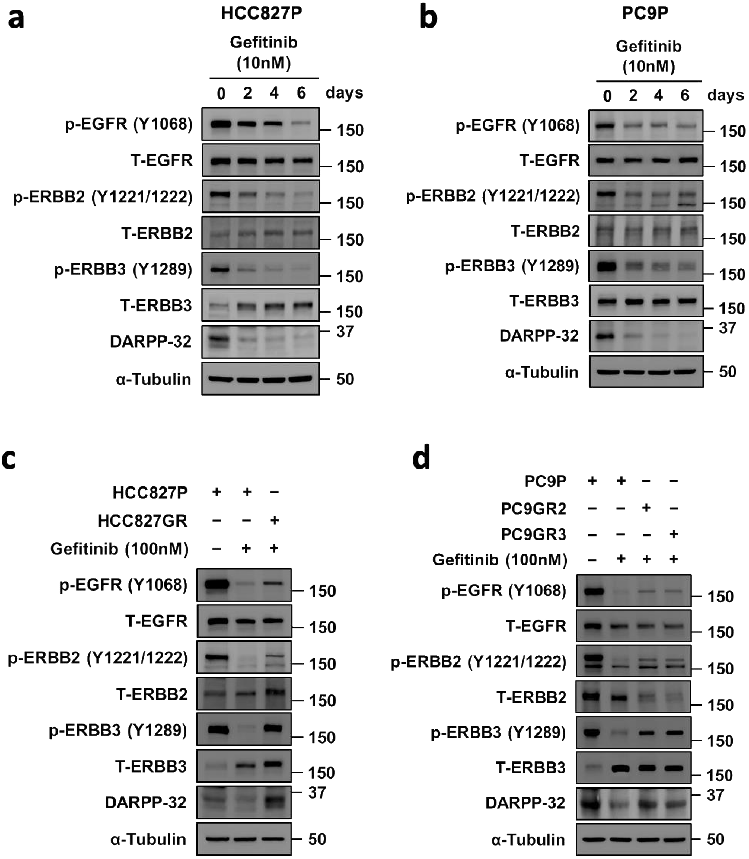
DARPP-32 is upregulated in gefitinib-resistant cells. a-b Human NSCLC HCC827P (a) and PC9P (b) cells treated with 10nM gefitinib at indicated days were immunoblotted with antibodies against phosphorylated EGFR (p-EGFR), total EGFR (T-EGFR), phosphorylated ERBB2 (p-ERBB2), total ERBB2 (T-ERBB2), phosphorylated ERBB3 (p-ERBB3), total ERBB3 (T-ERBB3), DARPP-32, and α-tubulin (loading control). c-d HCC827P, HCC827GR (c), PC9P, PC9GR2, and PC9GR3 (d) cells treated with 100nM gefitinib were lysed and antibody-reactive protein bands were detected using anti-p-EGFR, EGFR, p-ERBB2, ERBB2, p-ERBB3, ERBB3, DARPP-32, and α-tubulin antibodies. Immunoblotting experiments were repeated independently at least three times, and a representative experimental result is shown.

### DARPP-32 overexpression is associated with decreased EGFR TKI-induced NSCLC cell death

Given that DARPP-32 is upregulated in gefitinib-resistant NSCLC cells, we designed experiments to assess the functional effects of DARPP-32 overexpression in the presence of EGFR TKI. We stably silenced DARPP-32 protein expression in HCC827GR cells via lentiviral-mediated transduction of two previously validated shRNAs^28, 30^ (Supplementary Fig. 3a) and then analyzed cell viability in the presence of increasing gefitinib concentrations (Supplementary Fig. 3b). As evident by IC_50_ values, depletion of DARPP-32 decreases cell viability in HCC827GR cells upon gefitinib treatment (Supplementary Fig. 3b-c). We next stably overexpressed DARPP-32 in HCC827P cells (Supplementary Fig. 3d) and measured cell viability upon incubation with increasing concentrations of gefitinib (Supplementary Fig. 3e). HCC827P cells overexpressing DARPP-32 isoforms exhibit a greater IC_50_ value relative to controls (Supplementary Fig. 3e-f), suggesting that overexpression of DARPP-32 promotes resistance to gefitinib. We next sought to understand how DARPP-32 isoforms regulate EGFR-mutated NSCLC cell survival in the presence of gefitinib. We performed flow cytometry-based annexin V apoptosis studies in HCC827GR cells treated with increasing concentrations of gefitinib. We observed increased apoptotic cell death in DARPP-32-depleted HCC827GR cells upon gefitinib treatment compared to LacZ shRNA-transduced control cells (Fig. 2a; Supplementary Fig. 4). We next measured the expression of Poly (ADP-ribose) polymerase 1 (PARP1) and caspase-3 proteins in gefitinib-resistant, DARPP-32-silenced HCC827 and PC9 cells by western blot analysis. PARP-I and caspase-3 produce specific proteolytic cleavage fragments that are well-established surrogates of apoptotic cellular death^52^. We observed an increase in the expression of PARP-I and caspase-3 cleavage fragments in gefitinib-treated, DARPP-32-ablated cells relative to controls (Fig. 2b; Supplementary Fig. 5a-b), suggesting an anti-apoptotic role of DARPP-32. To validate that DARPP-32 overexpression does indeed promote cell survival, we performed annexin V assays. Indeed, our data suggests that stable overexpression of DARPP-32 isoforms protects HCC827P cells from gefitinib-induced apoptosis (Fig. 2c; Supplementary Fig. 6). Correspondingly, we observed a decrease in PARP-I and caspase-3 cleavage, suggesting overexpression of DARPP-32 isoforms in HCC827P and PC9P cells reduces gefitinib-mediated apoptotic cell death (Fig. 2d; Supplementary Fig. 5c). Collectively, our findings suggest DARPP-32 reduces gefitinib-induced apoptosis of EGFR-mutated NSCLC cells.

**Figure 2:**
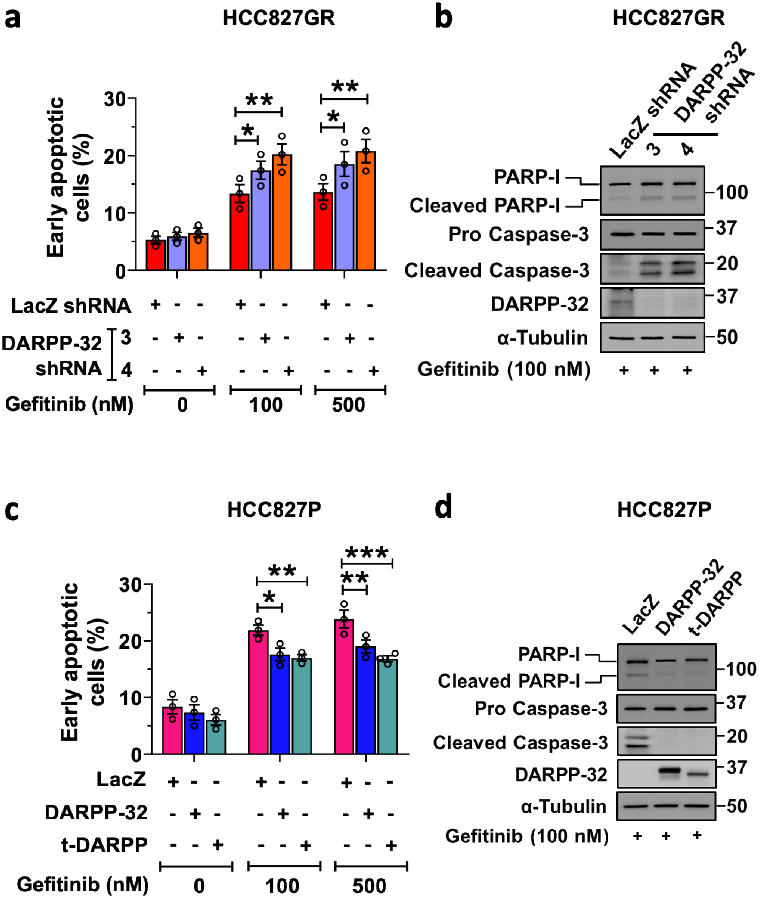
Overexpression of DARPP-32 represses gefitinib-induced cell apoptosis. **a** Lentivirus encoding control (LacZ) or DARPP-32 shRNAs were transduced in HCC827GR cells. Cells treated with gefitinib were used to measure apoptosis using FITC-conjugated anti-annexin V antibodies. **b** Immunoblotting was performed in lysates from HCC827GR cells transduced with LacZ or DARPP-32 shRNAs using antibodies against cleaved and uncleaved PARP-I, cleaved and uncleaved (i.e., pro-) caspase-3, DARPP-32 and α-tubulin (loading control). **c** Human NSCLC HCC827P cells were transduced with retrovirus containing control-(LacZ), DARPP-32- or t-DARPP-overexpressing clones. Flow cytometry-based apoptosis analysis was performed in gefitinib-treated cells to detect annexin V-positive cells. **d** HCC827P cells overexpressing LacZ, DARPP-32, or t-DARPP were lysed and separated in SDS-PAGE. Immune-reactive protein bands were detected using anti-PARP-I, caspase-3, DARPP-32, and α-tubulin antibodies. Each open circle on a graph represents an independent experiment. Immunoblot experiments were repeated at least three times. The average number of annexin V-positive cells of three independent experiments were plotted in a bar graph. Error bars indicate standard error of mean (SEM; n=3). *P<0.05, **P<0.01, and ***P<0.001, 2-way ANOVA followed by Dunnett’s test for multiple comparison.

### DARPP-32 upregulation and increased ERBB3 activation correlate with EGFR TKI resistance

We next sought to determine the molecular basis of DARPP-32-mediated cell survival in the presence of EGFR inhibition. DARPP-32 has been shown to promote resistance of gastric cancer cells to EGFR inhibitors by promoting an interaction between EGFR and ERBB3, which drives PI3K-AKT signaling^53^ to “bypass” EGFR TKI resistance. Importantly, we observe concomitant decreases in DARPP-32 protein expression and phosphorylation of EGFR, ERBB2, and ERBB3 over time when gefitinib-sensitive HCC827P and PC9P cells were treated with various doses of gefitinib (Fig. 1a-b; Supplementary Fig. 1a-b). We next replicated these parental cell experiments relative to their gefitinib resistant counterparts in the presence of an intermediate dose of gefitinib. The gefitinib-induced decreases in DARPP-32, p-EGFR, and p-ERBB3 that were observed in gefitinib-sensitive HCC827 and PC9P cells do not occur in gefitinib-resistant cells; HCC827GR, PC9GR2, and PC9GR3 all express high levels of DARPP-32, p-EGFR, and p-ERBB3 upon treatment with 100 nM gefitinib (Fig. 1c-d). Total ERBB3 protein levels were markedly increased upon EGFR TKI treatment in parental cells and these high ERBB3 levels were maintained in gefitinib-treated HCC827GR, PC9GR2, and PC9GR3 cells (Fig. 1c-d). Changes observed in p-ERBB2 and total ERBB2 protein expression were less uniformly consistent across the two resistance models, HCC827 and PC9 (Fig. 1c-d), and ERBB4 protein was undetectable in these cells (data not shown). Taken together, our observations suggest that upregulation of p-ERBB3, total ERBB3 and total DARPP-32 protein levels positively correlate with an EGFR TKI resistance phenotype in EGFR-mutated NSCLC cells.

### DARPP-32 stimulates ERBB3 activation in EGFR TKI-treated NSCLC cells

Given that DARPP-32 is overexpressed and ERBB3 is activated during gefitinib treatment in EGFR-mutated NSCLC cells, we propose a mechanism of acquired resistance to EGFR TKIs in NSCLC, in which DARPP-32 mediates a switch from EGFR TKI-sensitive EGFR homodimers to TKI-resistant EGFR:ERBB3 heterodimers. This hypothesis is supported by findings showing that the physical association of EGFR and ERBB3 promotes resistance to gefitinib in NSCLC^54^. To test this hypothesis, we assessed the phosphorylation status of EGFR and ERBB3 by immunofluorescence studies in EGFR TKI-sensitive PC9P human NSCLC cells overexpressing DARPP-32 or t-DARPP upon gefitinib treatment. We observed a substantial reduction of p-EGFR intensity in gefitinib-treated LacZ-overexpressed control PC9P cells, whereas p-EGFR intensity in EGFR-mutated cells overexpressing DARPP-32 isoforms remained unchanged upon gefitinib treatment (Fig. 3a-b). Overexpression of DARPP-32 and t-DARPP promotes increased p-ERBB3 upon EGFR TKI treatment (Fig. 3a,c), suggesting DARPP-32 upregulation may be associated with increased activation of ERBB3 in the presence of EGFR TKIs. We next asked whether stable shRNA-mediated depletion of DARPP-32 in gefitinib-resistant PC9GR3 cells affects phosphorylation of ERBB3 and EGFR upon gefitinib treatment. We observed a decrease in p-ERBB3 expression in gefitinib-treated DARPP-32-ablated PC9GR3 cells, whereas changes of p-ERBB3 levels were not detectable in corresponding LacZ shRNA control PC9GR3 cells upon treatment with gefitinib (Supplementary Fig. 7a,c). Others have reported that p-EGFR is not responsive to gefitinib in PC9GR3 cells^44^. Correspondingly, knockdown of DARPP-32 in gefitinib-treated PC9GR3 cells did not affect p-EGFR levels (Supplementary Fig. 7a-b). Collectively, our results demonstrating changes in activation of ERBB3 upon DARPP-32 modulation in the presence of EGFR TKI support a model in which DARPP-32 contributes to ERBB3-driven “bypass signaling” to promote EGFR-mutated NSCLC cell survival.

**Figure 3:**
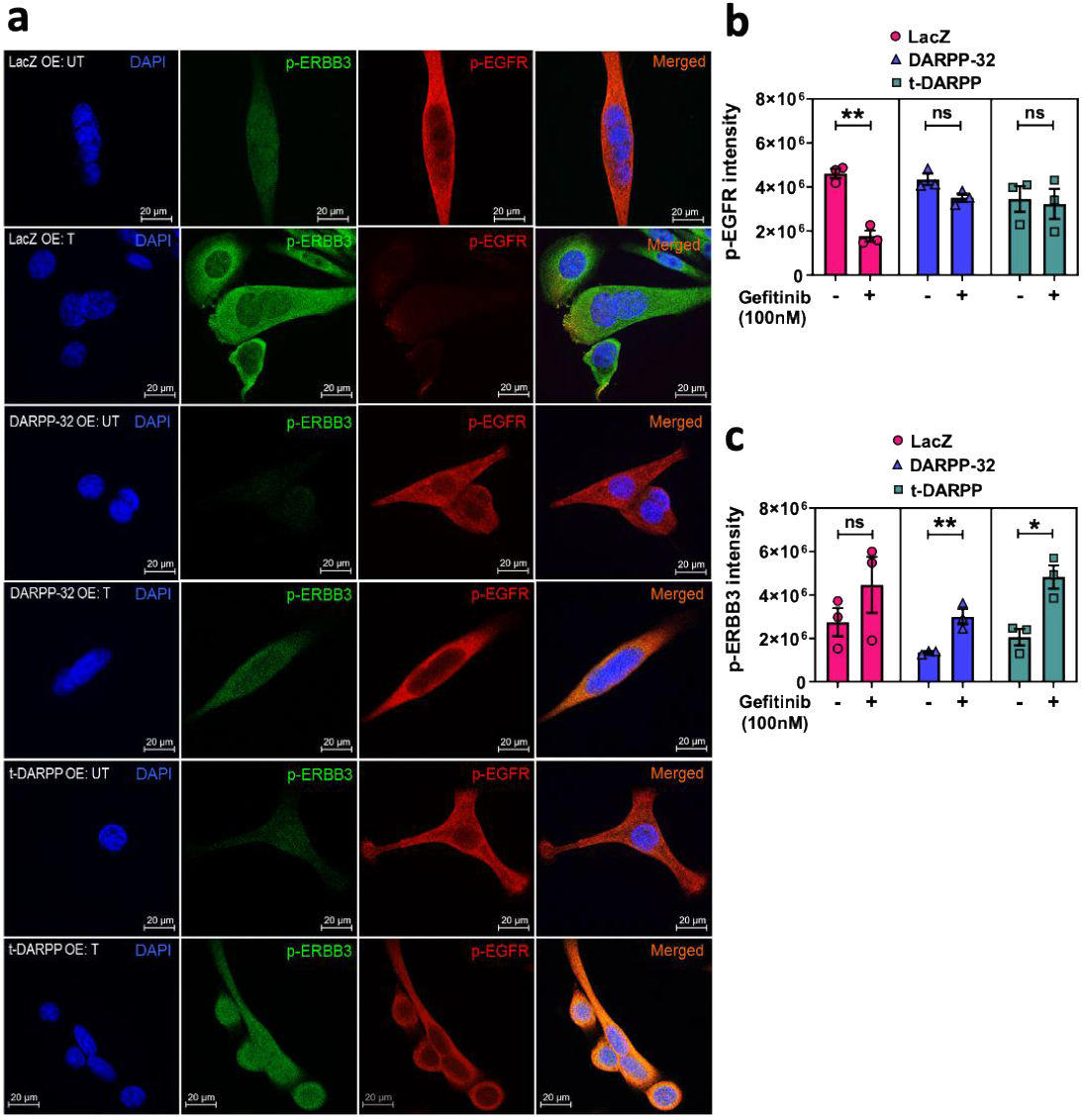
Overexpression of DARPP-32 isoforms increases p-ERBB3 expression. **a** Human lung cancer cell line PC9P treated with vehicle (UT) or 100nM gefitinib (T) were transduced with retrovirus containing control (LacZ)-, DARPP-32- or t-DARPP-overexpressing clones and immunofluorescence experiments were performed using primary antibodies against p-ERBB3 (green) and p-EGFR (red). Nuclei were stained with DAPI (blue). **b-c** Average red (b) and green (c) fluorescence intensity of 6-10 random microscopic fields for each sample was reported. Experiments were repeated at least three times. Scale bar, 20 μm. Bar graphs indicate mean ±SEM (n=3). *P<0.05 and **P<0.01, 2-way unpaired t-test.

### Physical association between EGFR and ERBB3 is positively regulated by DARPP-32 isoforms

To better understand the mechanism of resistance to gefitinib in EGFR-mutated NSCLC cells, we aimed to determine how DARPP-32 activates ERBB3 signaling to suppress gefitinib-mediated EGFR inhibition. To address our hypothesis that DARPP-32 drives EGFR:ERBB3 heterodimerization to evade EGFR TKI-mediated cell death, we performed immunoprecipitation studies to assess potential EGFR and ERBB3 interactions in DARPP-32-modulated EGFR-mutated NSCLC cells. Immunoprecipitation using anti-ERBB3 antibody demonstrates that EGFR and ERBB3 physically interact and that the EGFR:ERBB3 association increases upon overexpression of DARPP-32 isoforms in HCC827P and PC9P cells (Fig. 4a-b). Furthermore, immunoprecipitation for DARPP-32 in parental cells reveals that DARPP-32 physically interacts with EGFR and ERBB3 (Fig. 4a-b), suggesting it associates with the EGFR:ERBB3 complex. We sought to investigate how DARPP-32 affects EGFR and ERBB3 interactions in gefitinib-resistant cells relative to -sensitive parental cells. ERBB3 immunoprecipitation experiments suggest that DARPP-32 upregulation results in increased association of EGFR with ERBB3 in gefitinib-resistant HCC827GR, PC9GR2, and PC9GR3 cells relative to their gefitinib-sensitive counterparts (Fig. 4c-d). Upon DARPP-32 upregulation, we observed an increase in the association between DARPP-32 and ERBB3 in gefitinib-resistant cells with a concomitant decrease in EGFR:DARPP-32 association (Fig. 4c-d), suggesting DARPP-32 may promote EGFR:ERBB3 heterodimer formation in EGFR TKI-resistant cells.

**Figure 4:**
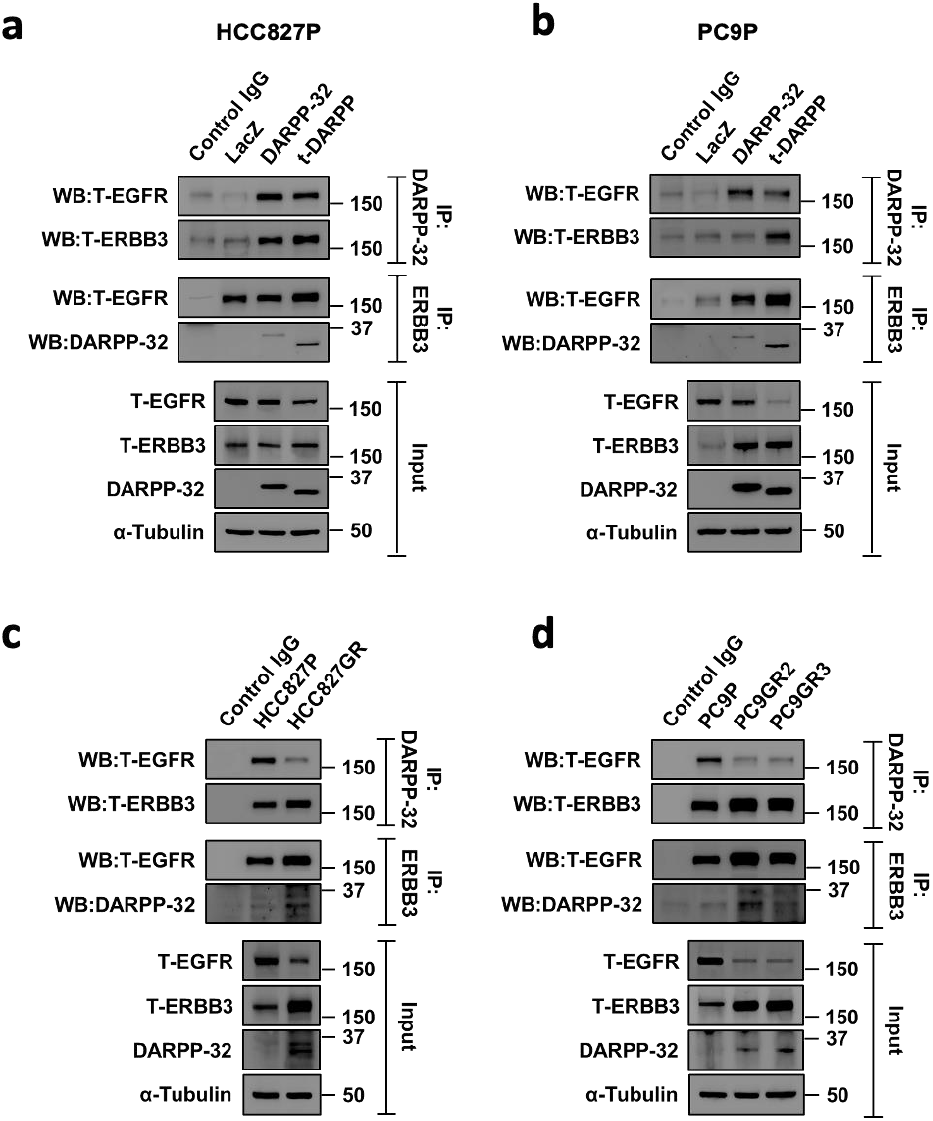
DARPP-32 physically associates with EGFR and ERBB3. **a-b** EGFR-mutated human NSCLC HCC827P (a) and PC9P (b) cells transduced with retrovirus encoding control (LacZ)-, DARPP-32- or t-DARPP-overexpressing clones were immunoprecipitated using antibodies against FLAG (detects both DARPP-32 isoforms), and ERBB3. Immunoprecipitated protein complexes and total cell lysates (input) were immunoblotted using anti-EGFR, ERBB3, FLAG, and α-tubulin antibodies. **c-d** Human lung adenocarcinoma HCC827P, HCC827GR (c), PC9P, PC9GR2, and PC9GR3 (d) cells were lysed and immunoprecipitated with anti-DARPP-32 (recognizes endogenous DARPP-32 and t-DARPP) and anti-ERBB3 antibodies. Immunoprecipitated lysates along with total cell lysates were separated on SDS-PAGE followed by immunoblot analysis using antibodies against EGFR, ERBB3, DARPP-32, and α-tubulin. Immunoprecipitation experiments were repeated at least three times.

### DARPP-32-driven p-EGFR/p-ERBB3 heterodimerization activates downstream MEK and PI3K/AKT/mTOR signaling

Given that ERBB3 has limited kinase activity and relies on heterodimerization with EGFR for activation^55^, we postulate that DARPP-32 promotes ERBB3 phosphorylation by increasing physical association between p-EGFR and p-ERBB3. To address our theory, we performed proximity ligation assay (PLA) using anti-p-EGFR and anti-p-ERBB3 antibodies in PC9GR3 cells. PLA is a powerful tool for identifying protein-protein interaction *in situ* with high specificity and sensitivity. Our PLA findings suggest that gefitinib treatment induces p-EGFR/p-ERBB3 heterodimer complex formation in PC9GR3 cells (Fig. 5a-b). However, ablation of DARPP-32 in PC9GR3 cells abolishes gefitinib-induced p-EGFR/p-ERBB3 dimerization, suggesting DARPP-32 plays a significant role in the formation of these active heterodimers (Fig. 5a-b). We next sought to determine how DARPP-32 regulates MEK/ERK and PI3K/AKT signaling pathways in the presence of gefitinib. It has been reported that ligand-independent EGFR activation initiates intracellular signaling via Ras/Raf/MEK/ERK and PI3K/AKT signaling pathways^13^. We show by immunoblotting that overexpression of DARPP-32 isoforms increases p-AKT and p-ERK expression in gefitinib-treated sensitive cells (Fig. 6a). Knockdown of DARPP-32 reduces p-AKT and p-ERK expression in gefitinib-treated resistant cells (Fig. 6b). EGFR-dependent PI3K activation requires dimerization with the ERBB3 receptor because docking sites of PI3K (i.e. p85 subunit) are abundant on ERBB3 and absent within EGFR^13^. To test our hypothesis that DARPP-32 activates the PI3K signaling pathway in EGFR-mutated NSCLC cells, we used a bioinformatics approach to assess DARPP-32 transcript expression in specimens derived from 80 EGFR-mutated NSCLC patients cataloged in The Cancer Genome Atlas (TCGA). We first subdivided patient-derived specimens into two groups based on high versus low DARPP-32 mRNA expression (Supplementary Fig. 8a). Interestingly, we found that expression of RPS6KB2 transcripts, but not RPS6KB1 transcripts, increases in lung tumor specimens with high expression of DARPP-32 (Supplementary Fig. 8b-c). The ribosomal S6 kinase isoforms (i.e. RPS6KB1, RPS6KB2) are downstream targets of PI3K/AKT/mTOR signaling^56^. Given that both kinases phosphorylate the 40S ribosomal protein S6^56^, we next assessed the expression of phosphorylated RPS6 proteins to determine whether increased expression of RPS6KB2 affects the phosphorylation status of RPS6. Our results reveal increased expression of phospho-RPS6 proteins (i.e. pS235/S236 and pS240/S244) in the patient-derived specimens with high DARPP-32 transcript levels. The expression of unmodified RPS6 proteins among high versus low DARPP-32 transcript groups is unchanged, suggesting that upregulation of RPS6KB2 results in RPS6 protein activation (Supplementary Fig. 8d-f). Taken together, our findings suggest DARPP-32 promotes dimerization of active EGFR and ERBB3 receptors to stimulate PI3K/AKT/mTOR and MEK/ERK signaling in EGFR TKI refractory NSCLC progression.

**Figure 5:**
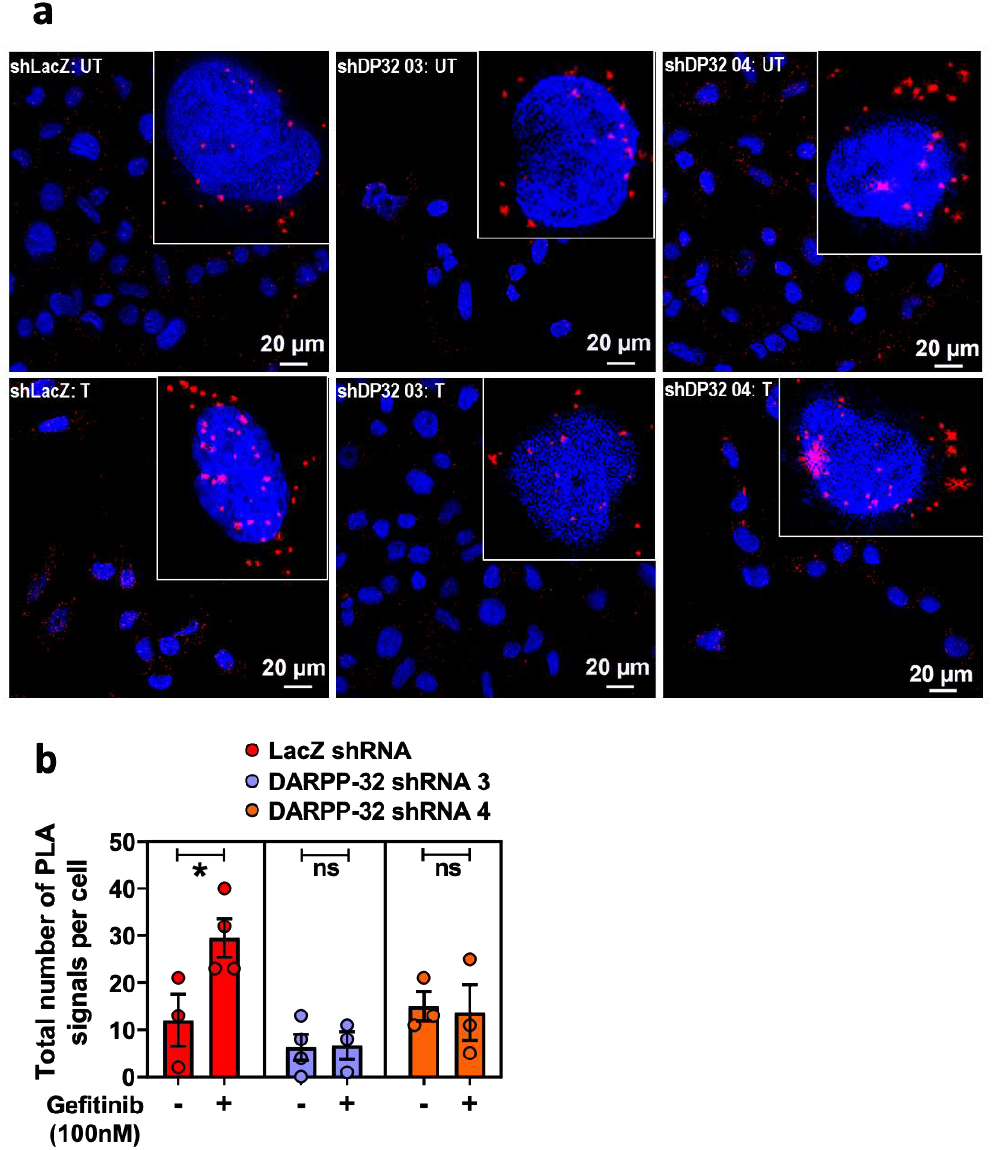
Depletion of DARPP-32 reduces p-EGFR to p-ERBB3 heterodimer formation. **a** Proximity ligation assays (PLA) were performed in PC9GR3 cells stably transduced with control (shLacZ) or DARPP-32 shRNAs (shDP32) using antibodies against phosphorylated ERBB3 and EGFR following 24h incubation with either vehicle (UT) or 100 nM gefitinib (T). The images show a maximum intensity projection of the raw image based on 10 z-planes. PLA signals are shown in red and the DAPI-stained nuclei in blue. Scale bar, 20 μm. **b** Total number of PLA signals per cell were reported after calculating red fluorescence signals of 6-10 random microscopic fields for each group. Each circle on a graph represents an independent experiment. Bar graphs represent mean ±SEM of three independent experiments. *P<0.05, 2-way unpaired t-test.

**Figure 6:**
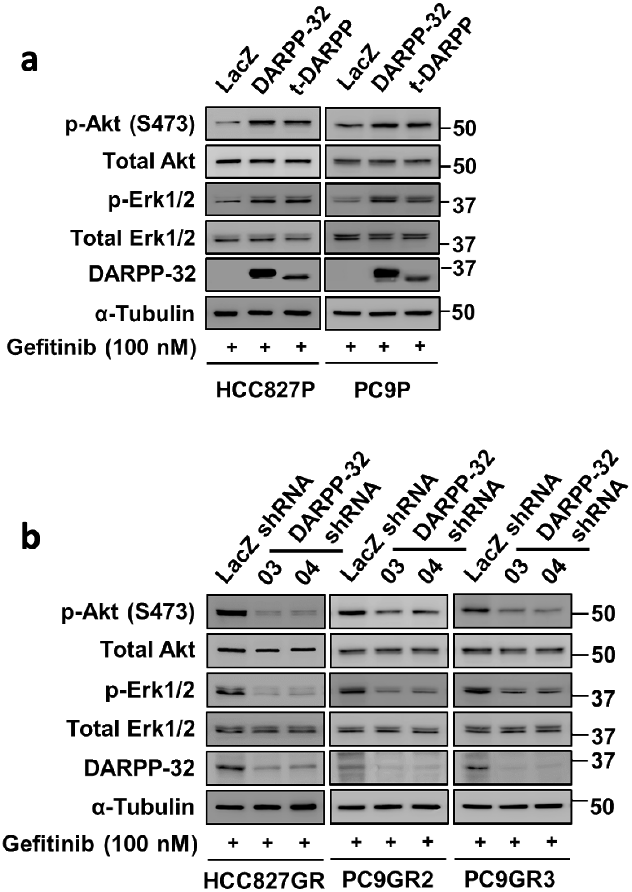
DARPP-32 activates AKT and ERK signaling in the presence of gefitinib. **a** Immunoblotting was performed in gefitinib-treated HCC827P and PC9P cells stably overexpressing control (LacZ), DARPP-32 or t-DARPP using antibodies against phosphorylated AKT (p-Akt), total AKT (Akt), phosphorylated ERK (p-Erk1/2), total ERK (Erk1/2), DARPP-32, and α-tubulin (loading control). **b** Gefitinib-resistant human lung cancer cell lines, HCC827GR, PC9GR2, and PC9GR3, were transduced with lentivirus containing LacZ or DARPP-32 shRNAs and treated with 100nM gefitinib for 24h. Cell lysates were separated and antibody-reactive protein bands were detected using anti-p-AKT, AKT, p-ERK, ERK, DARPP-32, and α-tubulin antibodies. Three independent immunoblotting experiments have been performed and representative results from one experiment have been shown.

### DARPP-32 promotes EGFR TKI refractory tumor growth *in vivo*

Based on our findings suggesting that DARPP-32 increases ERBB3 phosphorylation to bypass gefitinib-induced EGFR inhibition, we next sought to understand whether DARPP-32 drives NSCLC resistance to EGFR TKIs *in vivo*. To this end, we tested whether DARPP-32 ablation increases EGFR TKI sensitivity in a gefitinib-resistant orthotopic xenograft mouse model. Briefly, we injected luciferase-labeled human gefitinib-resistant (HCC827GR) NSCLC cells into the left thorax of anesthetized SCID mice, confirmed establishment of lung tumors via luciferase imaging, administered gefitinib over the course of two weeks, and measured tumors through non-invasive luciferase imaging (Fig. 7a). Mice challenged with HCC827GR cells with DARPP-32 stably silenced by shRNA show decreased tumor growth when treated every other day with gefitinib relative to vehicle controls (Fig. 7b; Supplementary Fig. 9a). DARPP-32 knockdown sensitizes gefitinib-resistant NSCLC tumors to EGFR inhibition *in vivo,* whereas no such effect was observed in mice challenged with control LacZ shRNA transduced HCC827GR cells (Fig. 7b; Supplementary Fig. 9a). Histological sections from these mice were immunostained for Ki-67. We observed decreased tumor cell proliferation in the lungs of gefitinib-treated mice challenged with DARPP-32-silenced HCC287GR cells (Fig. 7c), confirming that DARPP-32 knockdown enhances EGFR TKI-induced anti-cancer effects in gefitinib-resistant tumors *in vivo*.

**Figure 7:**
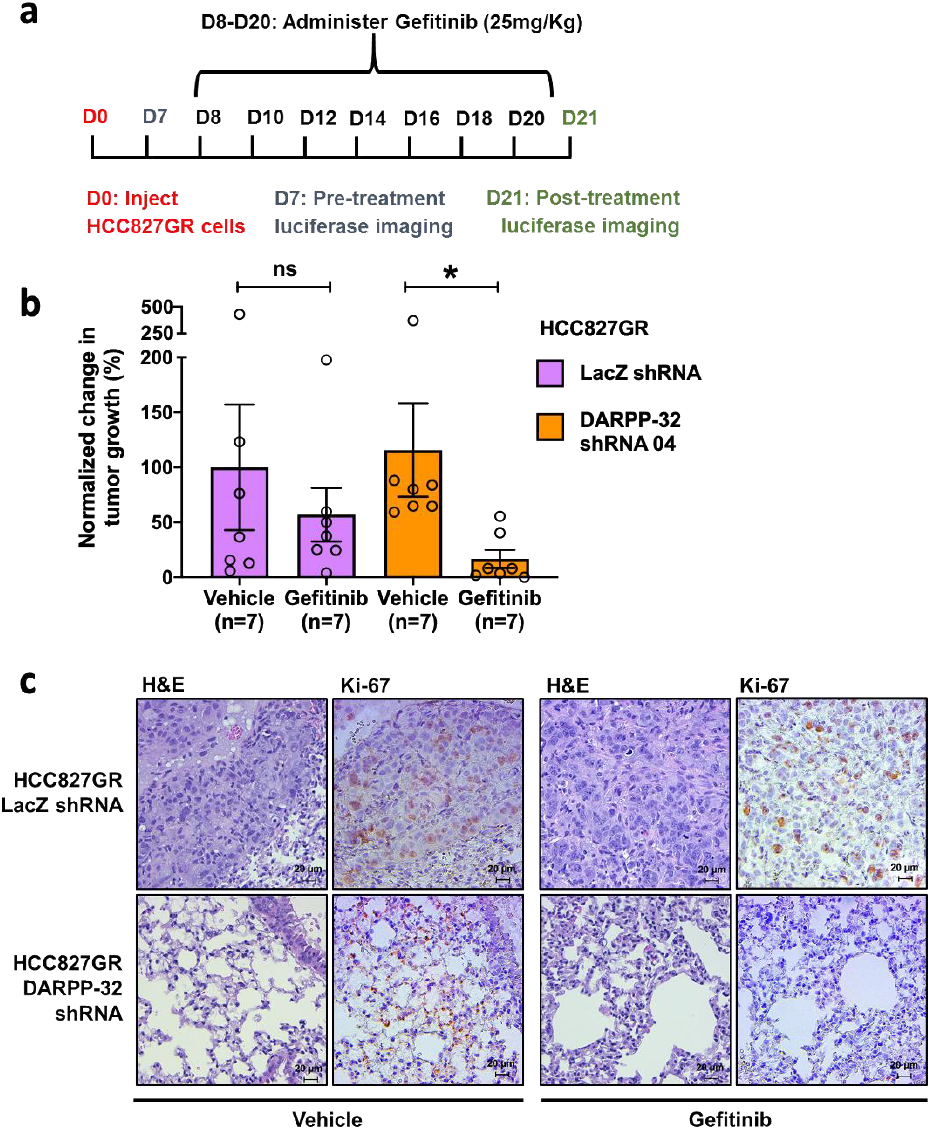
DARPP-32 silencing inhibits EGFR TKI refractory tumor growth *in vivo* **a** Luciferase-labeled human HCC827GR cells transduced with control (LacZ) or DARPP-32 shRNA were injected into the left thorax of SCID mice (n=7 mice per group), imaged for luminescence, administered 25mg/Kg gefitinib on indicated days. **b** Quantification of tumor growth was reported by determining the difference in relative luciferase units (RLU) before and after drug treatment. **c** Immunohistochemistry was performed using monoclonal Ki-67 antibody on formalin-fixed, paraffin-embedded lung tissue (n=3 mice per group) obtained from human lung tumor xenograft model. For evaluation of the morphology, slides were stained with hematoxylin and eosin (H&E) dye. Scale bar, 20 μm.

We next sought to determine whether overexpression of DARPP-32 isoforms promotes resistance to gefitinib *in vivo*. Gefitinib-sensitive HCC827 parental (HCC827P) tumors overexpressing DARPP-32 or t-DARPP that were implanted orthotopically into the lungs of mice exhibit gefitinib resistance relative to controls (Fig.8a; Supplementary Fig. 9b). To confirm the role of DARPP-32 in EGFR TKI resistance using a different NSCLC xenograft model, gefitinib-sensitive PC9P cells stably overexpressing DARPP-32 were subcutaneously injected into the flank of SCID mice. Once tumors reached ~150 mm^3^, mice were treated with EGFR TKI, gefitinib. Mice harboring DARPP-32 overexpressing PC9P tumors do not respond to gefitinib (Fig. 8b; Supplementary Fig. 10a), suggesting that DARPP-32 promotes resistance to gefitinib. Extirpated tumor volume and weight measurements confirm this finding (Supplementary Fig. 10b-c). Collectively, our observations demonstrate that DARPP-32 promotes gefitinib-refractory tumor growth *in vivo*. Taken together with our *in vitro* findings, our results indicate that DARPP-32 isoforms promote EGFR TKI-refractory NSCLC cell survival by stimulating the formation of active EGFR and ERBB3 heterodimers, increased AKT and ERK activation, and evasion of EGFR TKI-dependent apoptosis.

**Figure 8:**
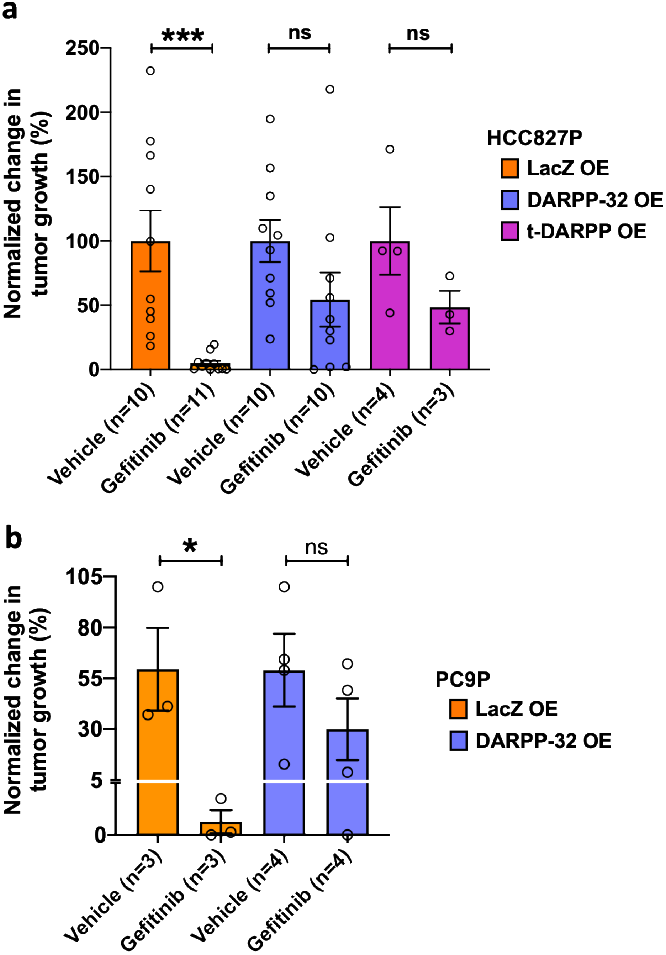
DARPP-32 overexpression protects EGFR TKI-sensitive human lung tumor xenografts from gefitinib-induced tumor reduction. **a** SCID mice were orthotopically injected with luciferase-labeled human HCC827P cells stably overexpressing control (LacZ), DARPP-32 or t-DARPP cDNAs. Vehicle- and gefitinib-treated mice were imaged for luminescence and quantification of HCC827P tumor growth before and after treatment was reported. Percentage change in tumor growth of 2 independent experiments were reported. **b** Human lung cancer PC9P cells transduced with retrovirus containing cDNA plasmids designed to overexpress control (LacZ) or DARPP-32 proteins were injected subcutaneously into the right flank of SCID mice. Mice were administered either vehicle or gefitinib (25mg/Kg) via oral gavage three times in a week once mean tumor volume reached 150 mm^3^. Tumor growth was recorded by measuring tumor volume with calipers before and after drug treatment. Experiments were concluded before average tumor volume exceeded 1500 mm^3^. Bar graphs represent mean ±SEM. Number of mice (n) used in each group was reported in the bottom of bar graphs. *P<0.05, ***P<0.001, 2-way unpaired t-test.

### Elevated DARPP-32 expression is associated with EGFR TKI resistance in NSCLC patients

To investigate the clinical relevance of DARPP-32 given its role in promoting resistance to EGFR first-generation TKIs in mouse models of human NSCLC, we assessed DARPP-32, p-EGFR, total EGFR, p-ERBB3, and total ERBB3 protein expression by immunostaining in paired EGFR TKI-naïve and -resistant specimens from 30 lung adenocarcinoma patients (Supplementary Table 1). Briefly, lung tumor specimens were biopsied from lung adenocarcinoma patients before EGFR TKI treatment (i.e. baseline) and following the development of progressive disease after first-line gefitinib or erlotinib therapy. For immunostaining of each protein, three pathologists independently scored the percentage of tumor cells staining positive and corresponding staining intensity (i.e., 0 = none, 1 = weak, 2 = moderate, 3 = strong expression). We calculated an immune reactive (IR) score for each specimen based on the percentage of tumor cells staining positive and the staining intensity in those cells (IR score = percentage of tumor cells x staining intensity). We found that DARPP-32, kinase-activated EGFR, total EGFR, kinase-activated ERBB3, and total ERBB3 proteins are upregulated in 1st generation EGFR TKI-resistant NSCLC patient-derived specimens relative to individual patient-matched (i.e. paired) baseline samples biopsied prior to frontline gefitinib or erlotinib treatment (Fig 9a-b). Collectively, our results suggest that DARPP-32 overexpression and increased EGFR and ERBB3 activation is associated with EGFR TKI resistance in NSCLC patients.

**Figure 9:**
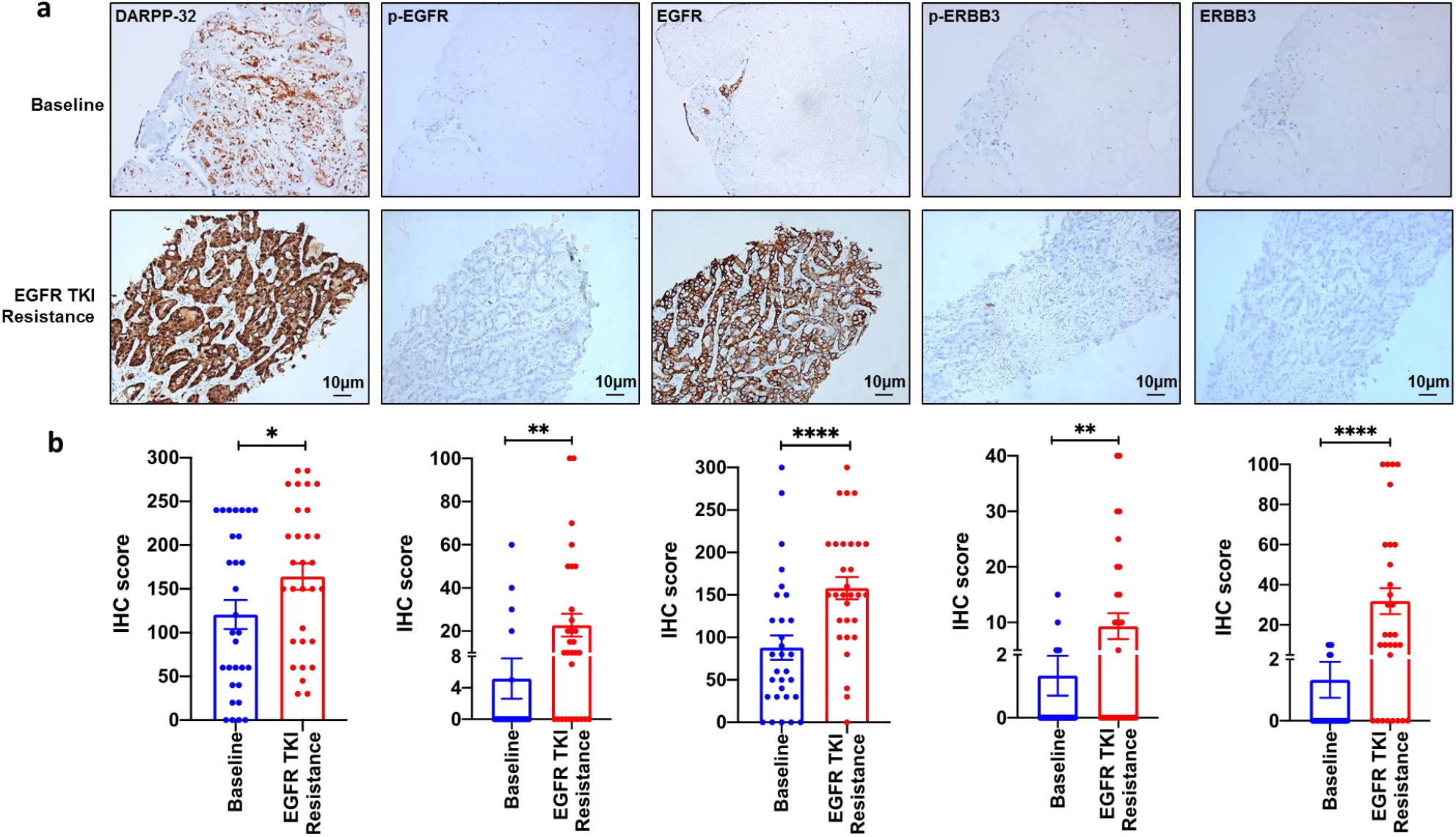
Expression of DARPP-32, p-EGFR, and p-ERBB3 proteins is elevated in EGFR TKI resistant lung adenocarcinoma. **a** Tumor tissue was biopsied before EGFR TKI treatment (i.e. baseline) and following EGFR TKI resistance (i.e. progressive disease after first-line gefitinib or erlotinib therapy) from lung adenocarcinoma patients with EGFR activating mutations (n=30 patients in each group). Paired baseline (top) and EGFR TKI resistance (bottom) lung tumor specimens were immunostained for DARPP-32, phosphorylated EGFR (p-EGFR), total EGFR, p-ERBB3, and total ERBB3. **b** IHC score was calculated by multiplying the staining intensity score (0-3) by the percent of positive tumor cells. Each circle on the plots represents single patient. *P<0.05, **P<0.01, ****P<0.0001,2-way unpaired t-test

## Discussion

Lung cancer is the deadliest and most frequently diagnosed type of tumor worldwide, with 1.6 million deaths reported annually^57^. The molecular targeting of specific oncogenic drivers has emerged as a major advancement in the treatment of NSCLC. Patients diagnosed with advanced non-squamous cell NSCLC are tested for oncogenic alternations and treated accordingly^20, 58^. Single oncogenic driver mutations in EGFR that confer sensitivity to TKIs are the most common targetable molecular alteration in lung adenocarcinoma. Although EGFR mutation positive patients initially respond well to EGFR TKI therapy, most patients inevitably develop resistance and experience rapid advanced disease progression. Developing acquired resistance to lung cancer therapy is a major problem. The development of effective strategies to circumvent the emergence of this resistance is needed to improve survival rates and the quality of life of NSCLC patients.

The spectrum of identified EGFR resistance mechanisms includes on-target EGFR gatekeeper mutations (i.e. EGFR^T790M^), amplifications of MET and ERBB2, MAPK-PI3K signaling activation, cell cycle alterations, rearrangements of RET or ALK kinases, and various other genomic alternations^59, 60^. Here, we sought to better understand the molecular mechanisms that control aberrant ERBB family signaling that drives EGFR TKI refractory lung adenocarcinoma progression. ERBB receptor tyrosine kinase family members, including ERBB1-ERBB4 (also known as EGFR, HER2, HER3, and HER4), consist of a single hydrophobic transmembrane region flanked by an extracellular ligand-binding domain and an intercellular tyrosine kinase domain^14^. ERBB family member signaling activates interconnected pathways that promote oncogenesis, including phosphatidylinositol-3 kinase (PI3K)–AKT–mammalian target of rapamycin (mTOR) and mitogen-activated protein kinase (MAPK) signal transduction^61, 62^, as well as phospholipase Cγ (PLCγ)–protein kinase C (PKC)^63, 64^ and Janus kinase 2 (JAK2)–signal transducer and activator of transcription 3 (STAT3) pathways^65, 66^. ERBB3, specifically, has been implicated in the initiation of EGFR TKI resistance. Unlike its fellow family members, ERBB3 was initially believed to be an inactive kinase because its kinase domain lacks certain residues known to be essential for catalytic activity^67^. However, ERBB3 forms heterodimers with other ERBB family members to become transphosphorylated and transactivated to sustain transduction of downstream oncogenic signaling that would otherwise be inhibited by EGFR TKIs acting upon EGFR homodimers^68–70^. Several known mechanisms of ERBB3-induced TKI resistance exist by which ERBB3 compensates for TKI-inhibited EGFR to trigger and sustain PI3K/Akt signal transduction. First, MET amplification has been shown to result in constitutive activation of ERBB3 signaling to promote gefitinib resistance in lung cancer cell lines^43^. Second, ERBB3 heterodimerization with ERBB2 has been demonstrated to drive oncogenic signaling in breast cancer^71^ as the effects of ERBB2 inhibition could be reversed by increasing ERBB3 phosphorylation and activity to drive a TKI resistance phenotype^70^. Third, ligand-mediated activation of ERBB3 has been shown to result in PI3K/Akt-mediated resistance to TKIs in a variety of cancers, including ERBB2-amplified breast cancer cells stimulated with ERBB3 ligands, NRG1^72^ or HRG^73^. We identify a new mechanism of ERBB3-mediated TKI resistance in which DARPP-32 physically stimulates this process of EGFR:ERBB3 heterodimer formation to promote PI3K/Akt and MAPK signaling to overcome the inhibitory effects of EGFR TKIs.

We were the first to report that DARPP-32 overexpression in lung cancer contributes to oncogenic growth^30^. While DARPP-32 is virtually undetectable in normal human lung^28^, DARPP-32 is overexpressed in human EGFR-mutated NSCLC. Specifically, we previously demonstrated that DARPP-32 proteins promote NSCLC cell survival through increased Akt and Erk1/2 signaling^30^. Given that these PI3K and MAPK signaling pathways are upregulated during resistance, we hypothesized that overexpression of DARPP-32 proteins in EGFR-mutated NSCLC may promote EGFR:ERBB3 “bypass signaling” that enables tumor cells to evade EGFR TKI monotherapy. In this report, we provide the first evidence that DARPP-32 overexpression in EGFR-mutated lung adenocarcinoma promotes ERBB3-mediated oncogenic signaling to drive EGFR TKI therapy refractory cancer progression. *In vivo* studies reveal that ablation of DARPP-32 protein activity sensitizes gefitinib-resistant lung tumor xenografts to EGFR TKI treatment, while DARPP-32 overexpression increases gefitinib-refractory lung cancer progression in gefitinib-sensitive lung tumors orthotopically xenografted into mice. Findings from proximity ligation assays, immunoprecipitation studies, and immunofluorescence experiments presented here support a model in which DARPP-32 mediates a switch from EGFR TKI-sensitive EGFR homodimers to TKI-resistant EGFR:ERBB3 heterodimers to potentiate oncogenic AKT and ERK signaling that drives therapy refractory tumor cell survival. To our knowledge, no proteins have been identified that are capable of mediating such a “dimerization switch” in EGFR-mutated NSCLC. Here, we take advantage of a unique cohort of paired tumor specimens derived from 30 lung adenocarcinoma patients before and after the development of EGFR TKI refractory disease progression to reveal that DARPP-32 as well as kinase-activated EGFR and ERBB3 proteins are overexpressed upon acquired EGFR TKI resistance. This observation coincides with our published report that increased t-DARPP immunostaining positively correlates with increasing T stage among unknown EGFR mutation status NSCLC patients^30^. There is no precedent of DARPP-32 isoform immunostaining in molecular targeted therapy naïve vs. resistant patients in other tumor types.

Our data comprehensively suggests that DARPP-32 overexpression promotes EGFR TKI resistance by stimulating formation of EGFR:ERBB3 heterodimers, which are less sensitive to EGFR inhibition and drive oncogenic signaling. Therefore, dual inhibition of EGFR and ERBB3 may better prevent treatment-refractory cancer progression as opposed to solely targeting EGFR, especially in tumors overexpressing DARPP-32. A precision oncology approach could be used to identify EGFR-mutated lung adenocarcinomas with high DARPP-32 and phosphorylated ERBB3 expression with the highest likelihood to benefit from dual EGFR and ERBB3 inhibition. For example, duligotuzumab is a human IgG1 monoclonal “two-in-one” antibody with high affinity for EGFR (K_D_ ~ 1.9 nM) and ERBB3 (K_D_ ~ 0.4 nM) developed to improve treatment response of solid tumors exhibiting ERBB3-mediated resistance to EGFR-targeted treatment^74^. Partial responses to duligotuzumab were achieved in patients with squamous cell carcinoma of the head and neck that had become resistant to cetuximab, an antibody therapy that inhibits EGFR^75^. Efficacy was also observed in tumors refractory to both radiation and long-term EGFR-targeted treatment^76, 77^. Duligotuzumab monotherapy has been shown to be well-tolerated in patients with locally advanced or metastatic solid tumors of epithelial origin^75^. However, a recent randomized phase II study of duligotuzumab vs. cetuximab in squamous cell carcinoma of the head and neck (i.e. MEHGAN; NCT01577173) found duligotuzumab did not improve disease free survival compared to cetuximab^78^. However, poor patient selection may have confounded its outcome as suggested by Dr. Saba in an associated *Commentary*^79^. Regardless, dual inhibition of EGFR and ERBB3 warrants further clinical investigation in trials that focus specifically on EGFR treatment-resistant patient populations^80^. Such clinical trials have not been performed and nor have any duligotuzumab studies in the clinic focused on EGFR-mutated LUAD patients. Future studies evaluating a dual EGFR and ERBB3 inhibitory approach in models of acquired EGFR TKI resistance are warranted, given that DARPP-32-mediated, ERBB3-driven resistance is a key mechanism of EGFR TKI-refractory LUAD progression, combined with the well-established safety profile of duligotuzumab.

## Acknowledgements

This work was supported by the Elsa U. Pardee Foundation, Institutional Research Grant #129819-IRG-16-189-58-IRG81 from the American Cancer Society, and The Hormel Foundation (to L.H.H.) as well as the Fifth District Eagles Cancer Telethon Postdoctoral Fellowship Award (to S.K.A.). M.A.K. is an employee of the Minneapolis VA Healthcare System. This material is based partially upon work supported by the Department of Veterans Affairs (specifically, the Veterans Health Administration). We thank Dr. Wael El-Rifai, Dr. Debabrata Mukhopadhyay, Dr. Scott Lowe, Dr. Aaron Hata, and Dr. Pasi Jänne for generously sharing plasmids and cell lines. We are grateful to Dr. Aaron Mansfield for facilitating collaborations that made this work possible. We thank The Hormel Institute and its staff for administrative, shared equipment, animal facility, and institutional support.

## Author contributions

S.K.A. conducted in vitro cell line-based experiments, including immunoprecipitation experiments, immunofluorescence studies, proximity ligation assays, apoptosis assays, etc. S.K.A. and L.H.H. managed the mouse colony and performed tumor studies in mice. S.K.A., L.W., and C.E.H. conducted murine in vivo imaging and necropsy. S.K.A., L.W., and Z.Z. performed the human cell line-based subcutaneous xenograft mice study. S.K.A. and Z.Z. performed western blotting experiments. Y. Zhang led the collection, immunostaining, imaging, pathological review, and analysis of patient-derived lung tumor specimens. Y. Zhou assisted with imaging and analysis of immuonstained specimens. N.Y. acquired the patient-derived specimens and provided associated clinical annotation. J.L., X.C., and L.Z. performed the pathological evaluation of immunostained patient-derived specimens. M.A.K. and S.K.A. performed computational biology analysis of TCGA data. S.K.A, Y. Zhang, L.W., M.A.K., and L.H.H. provided technical and scientific support. S.K.A. and L.H.H. performed experimental troubleshooting, reviewed relevant scientific literature, critically analyzed data, prepared figures, and wrote the manuscript. L.H.H. conceived the aims, led the project, and acquired funding to complete the reported research.

## Disclosures

None

## Supplemental Material

Supplementary Figures 1-10, Supplementary Table 1

**Supplementary Figure 1:**
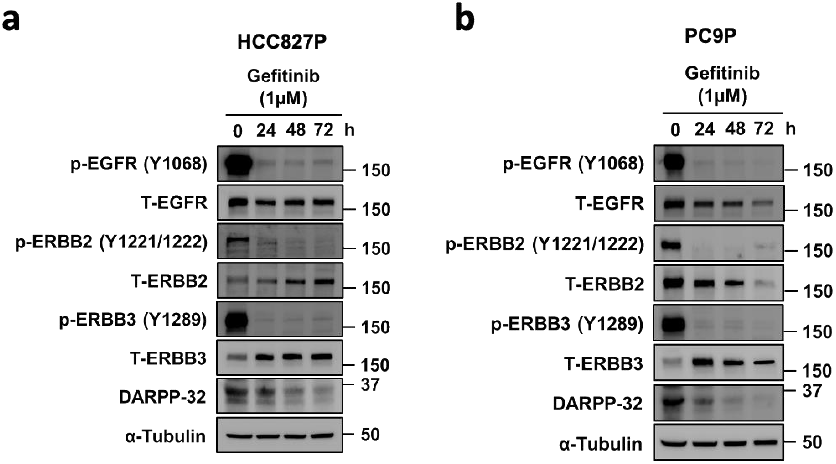
Gefitinib blocks EGFR phosphorylation in EGFR-mutated human NSCLC cells. **a-b** HCC827P (a) and PC9P (b) cells were lysed in RIPA buffer and phosphorylated EGFR (p-EGFR), total EGFR (T-EGFR), phosphorylated ERBB2 (p-ERBB2), total ERBB2 (T-ERBB2), phosphorylated ERBB3 (p-ERBB3), total ERBB3 (T-ERBB3), DARPP-32, and α-tubulin (loading control) proteins were detected by immunoblotting of cell lysates.

**Supplementary Figure 2:**
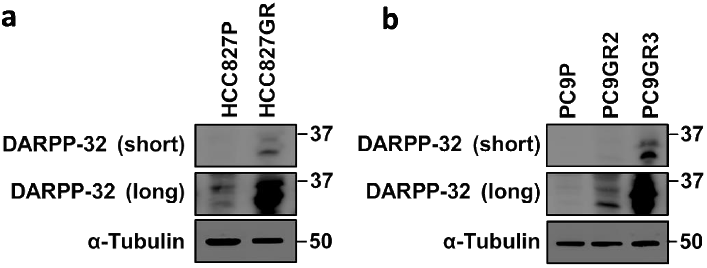
DARPP-32 is upregulated in gefitinib-resistant cell lines. **a** Human NSCLC cell lines, HCC827P and HCC827GR, were lysed and immunoblotted to detect DARPP-32 and α-tubulin (loading control) protein expression. **b** PC9P, PC9GR2, and PC9GR3 cell lysates were separated in SDS-PAGE and immunoblotting was performed using antibodies against DARPP-32 and α-tubulin. **a, b** Both “short” and “long” immunoblotting development exposures are depicted.

**Supplementary Figure 3:**
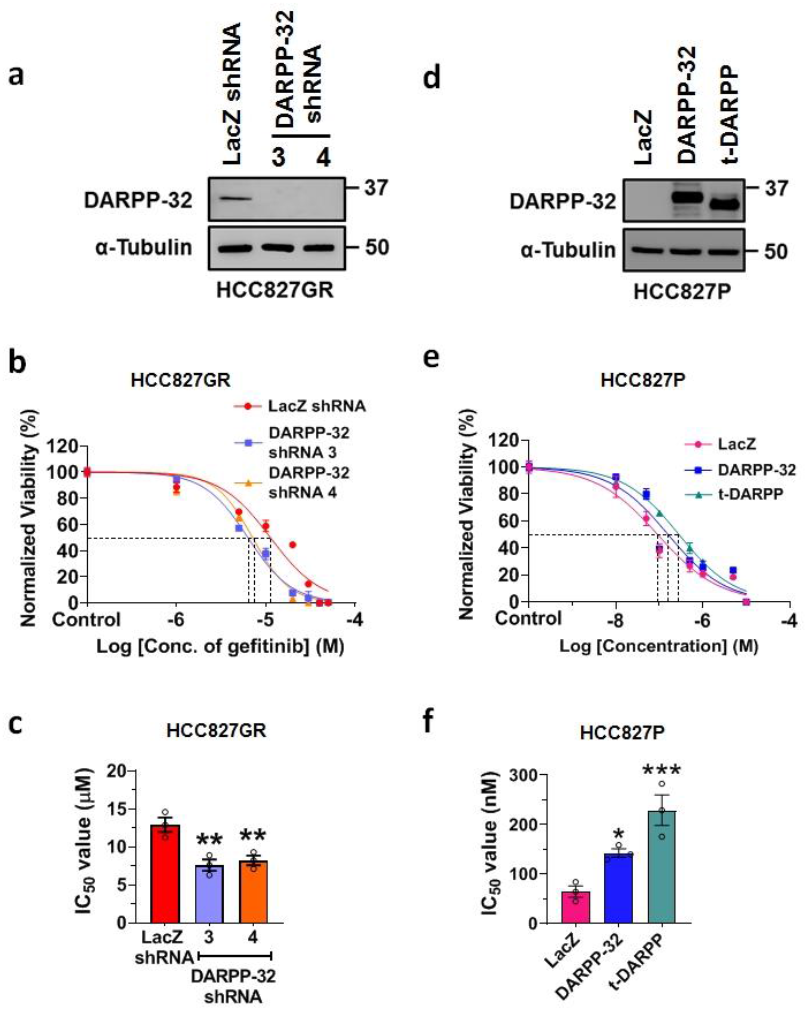
DARPP-32 overexpression increases NSCLC cell survival. **a** Human lung adenocarcinoma HCC827GR cells transduced with lentivirus encoding control (LacZ) or DARPP-32 shRNAs were immunoblotted with anti-DARPP-32 and -α-tubulin (loading control) antibodies. **b** HCC827GR cells were transduced with control (LacZ) or DARPP-32 shRNAs and seeded into 96-well cell culture plates. Cells were treated with increasing concentration of gefitinib and colorimeter-based cell survival assay was conducted using MTS1 reagents. **c** The half maximal inhibitory concentration (IC_50_) of gefitinib was determined from MTS1 survival assays and plotted. **d** HCC827P cells were transduced with retrovirus encoding control (LacZ), DARPP-32 or t-DARPP overexpressing clones. Cells were lysed and immunoblotting was performed to detect α-tubulin and DARPP-32 isoforms. **e** Cell survival assays were performed using HCC827P cells stably overexpressing LacZ, DARPP-32 or t-DARPP proteins exposed to increasing concentrations of gefitinib. **f** Gefitinib-treated HCC827P cells overexpressing LacZ or DARPP-32 isoforms were subjected to MTS1-based cell survival assays and IC_50_ of gefitinib was calculated. Each open circle on a graph represents an independent experiment. All bar graphs represent mean ± SEM (n=3). *P<0.05, **P<0.01, and ***P<0.001, one-way ANOVA followed by Dunnett’s test for multiple comparison.

**Supplementary Figure 4:**
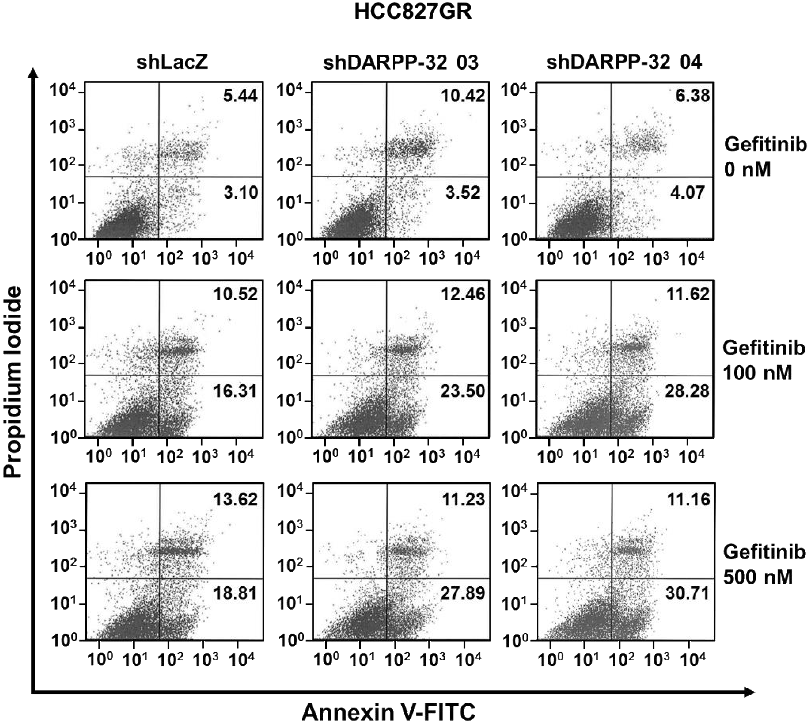
DARPP-32 ablation increases cell apoptosis in the presence of gefitinib. Human lung adenocarcinoma cell line, HCC827GR, transduced with control (LacZ) or DARPP-32 shRNAs were incubated with anti-annexin V antibodies conjugated with FITC followed by propidium iodide incorporation. The total number of annexin V-positive cells was determined using flow cytometry-based apoptosis assays. The numerical values on quadrants of the scatter plots represent the percentage of total cells in one single representative experiment.

**Supplementary Figure 5:**
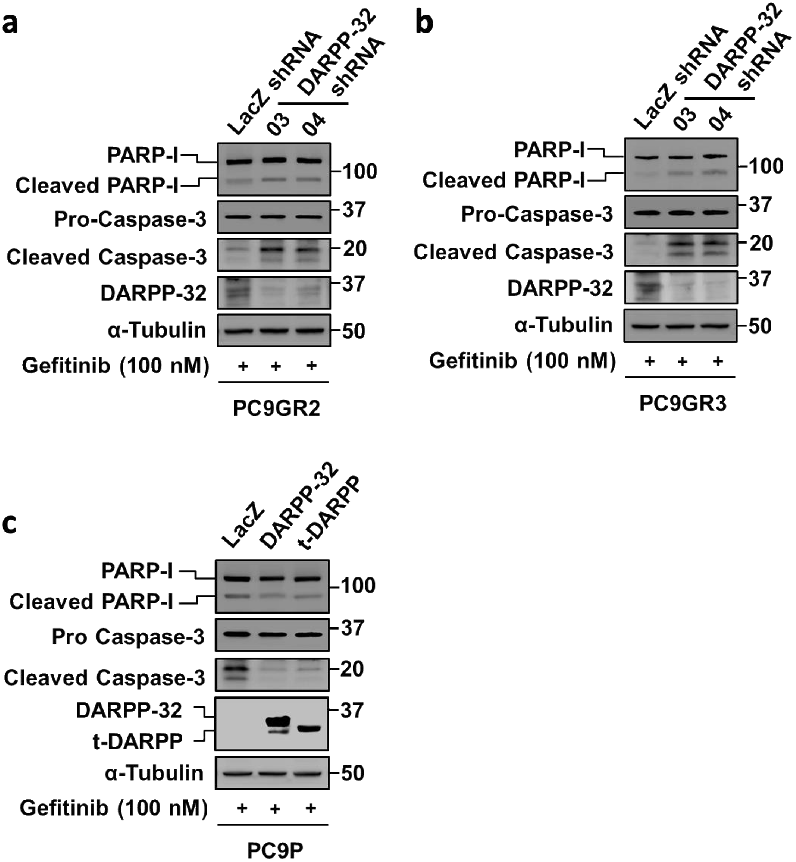
DARPP-32 depletion promotes gefitinib-induced cell death. **a-c** Gefitinib-treated DARPP-32-depleted PC9GR2 (a), and PC9GR3 (b) cells together with PC9P (c) cells overexpressing DARPP-32 isoforms were lysed and western blot was performed using antibodies against cleaved and uncleaved PARP-I, cleaved and uncleaved (i.e., pro-) caspase-3, DARPP-32 and α-tubulin (loading control).

**Supplementary Figure 6:**
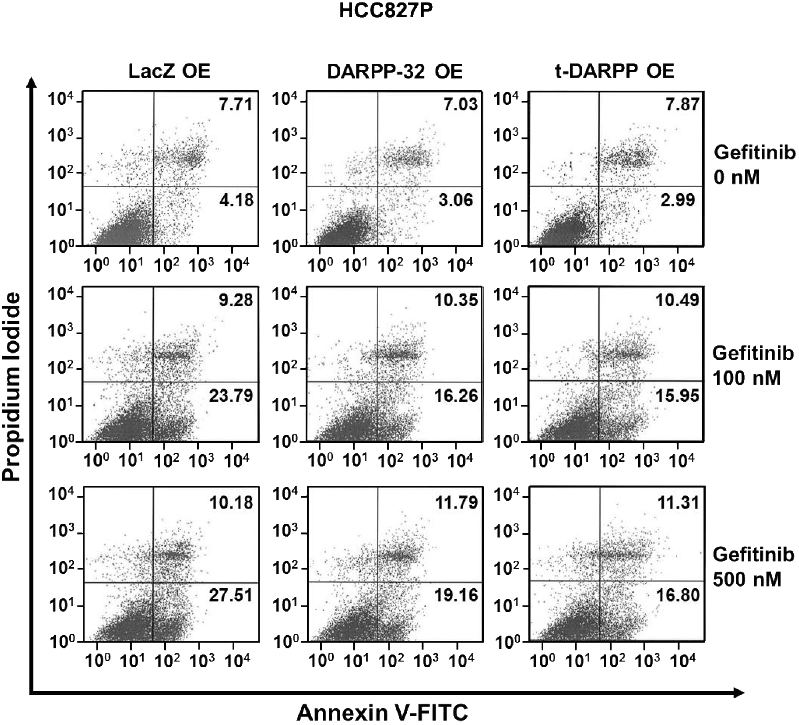
DARPP-32 overexpression suppresses cellular apoptosis upon gefitinib treatment. HCC827P cells were transduced with retrovirus containing control (LacZ)-, DARPP-32- or t-DARPP-overexpressing clones. Flow cytometry-based apoptosis assays were performed in gefitinib-treated cells using FITC-conjugated anti-annexin V antibodies along with propidium iodide. The numerical values on quadrants of the scatter plots represent the percentage of total cells in one single representative experiment.

**Supplementary Figure 7:**
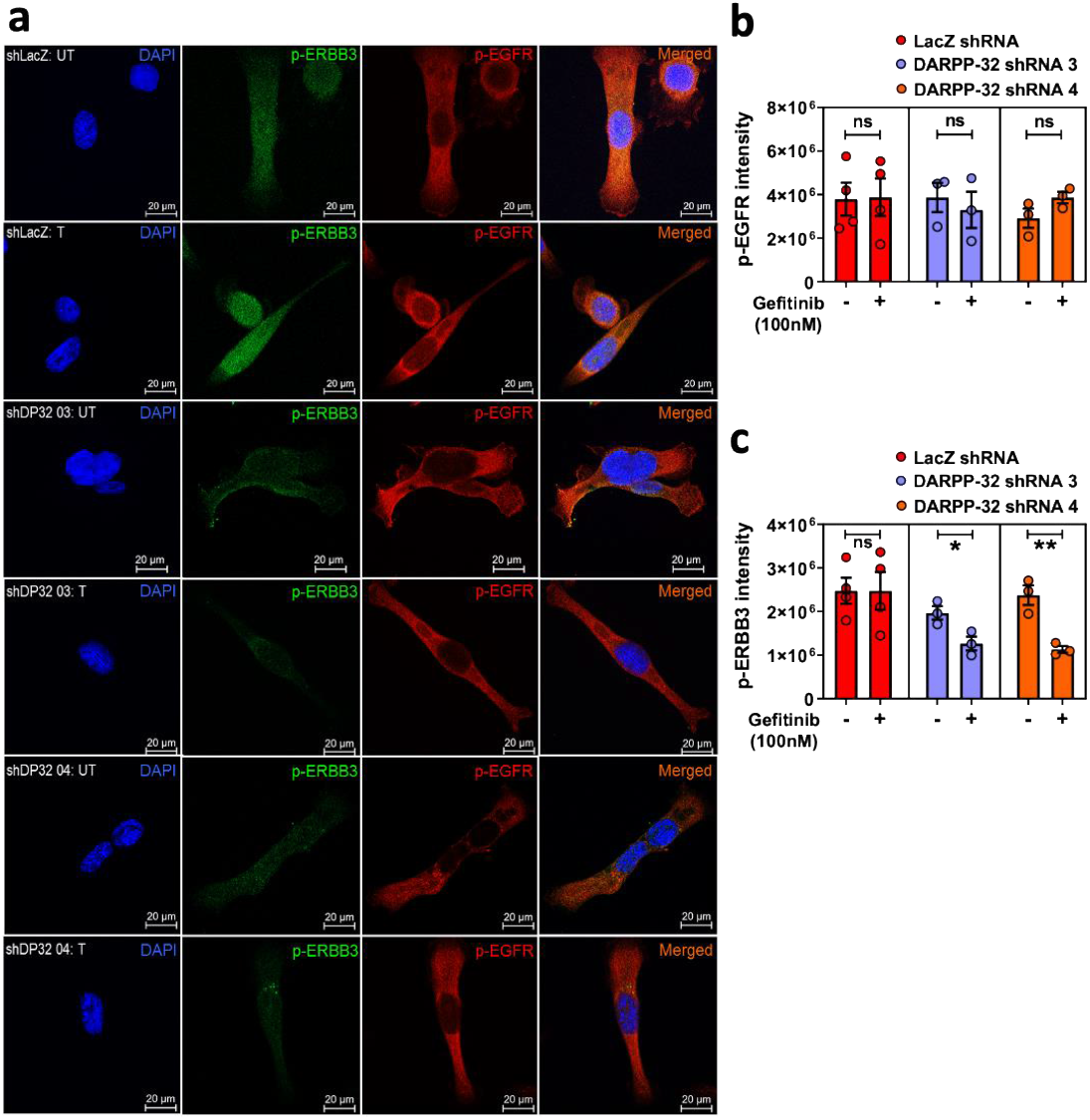
Expression of p-ERBB3 is controlled by DARPP-32. **a** PC9GR3 cells were transduced with lentivirus containing control (LacZ) or DARPP-32 shRNAs. Cells treated with vehicle (UT) or 100nM gefitinib (T) were fixed, permeabilized, and incubated with primary antibodies that detect p-ERBB3 (green) and p-EGFR (red) proteins. DAPI-stained nuclei were represented in blue color. **b-c** Expression of p-EGFR (b) and p-ERBB3 (c) was reported by calculating average fluorescence intensity of 6-10 random microscopic fields for each sample. Each circle on a graph represents an independent experiment. Scale bar, 20 μm. Results represent mean ±SEM (n=3). *P<0.05 and **P<0.01, 2-way unpaired t-test.

**Supplementary Figure 8:**
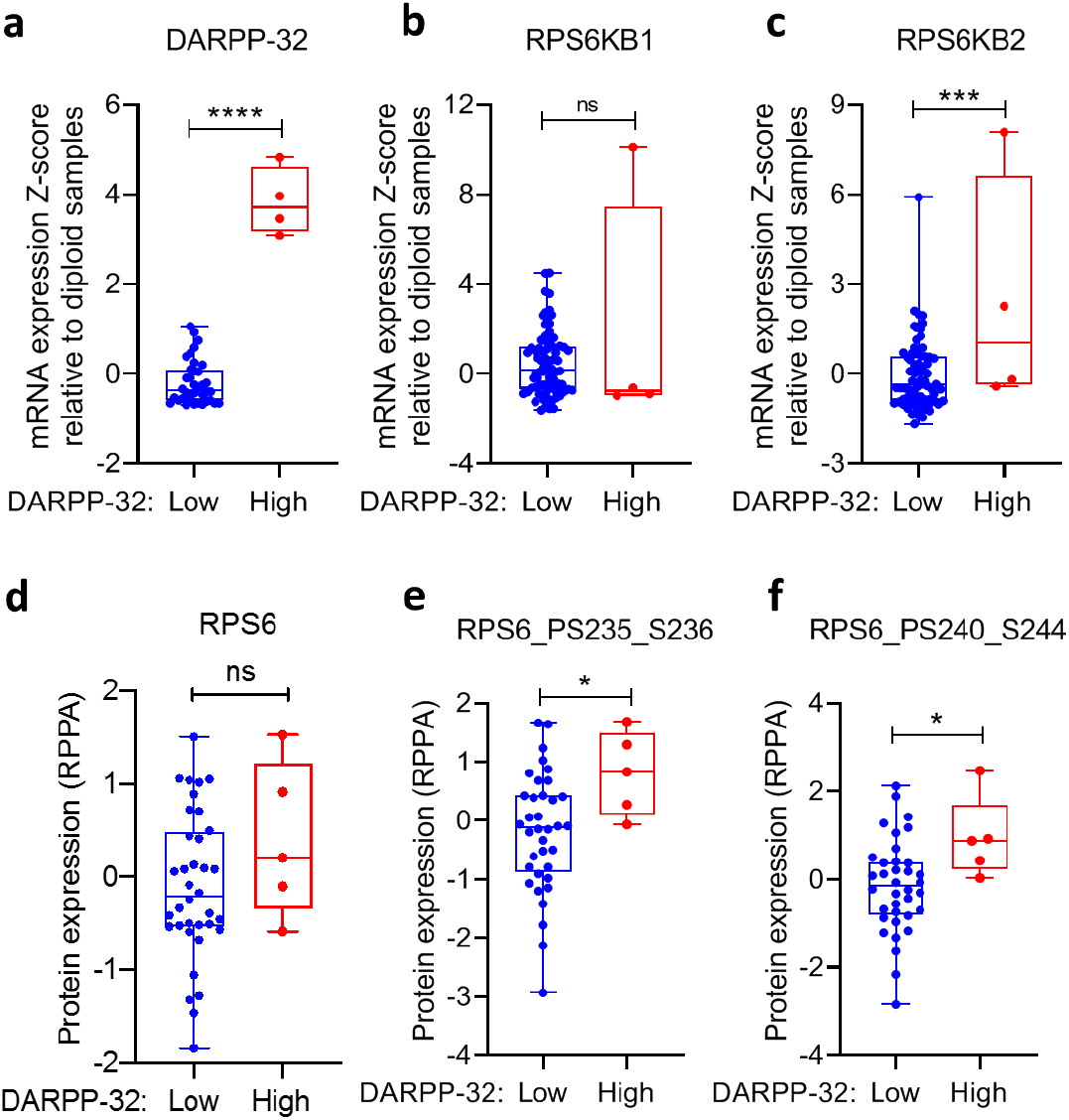
Integrative approach to analyze ribosomal protein S6 (RPS6) expression in human NSCLC patients using cBioPortal. **a-c** Box plot showing the relative mRNA expression of DARPP-32 (a), RPS6KB1 (b), and RPS6KB2 (c) genes in 80 human non-small cell lung adenocarcinoma patient samples from The Cancer Genome Atlas (TCGA) study. **d-f** Box plots representing the relative amount of total- (d) and phospho-RPS6 (e-f) proteins in DARPP-32-altered human NSCLC patients. Each patient has mutations in the EGFR gene. Based on the DARPP-32 expression, patients were divided between DARPP-32 -low (n=76) vs - high (n=4) groups. Each dot on box plots represents a single patient. *P<0.05, ***P<0.001, and ****P<0.0001, 2-way unpaired t-test.

**Supplementary Figure 9:**
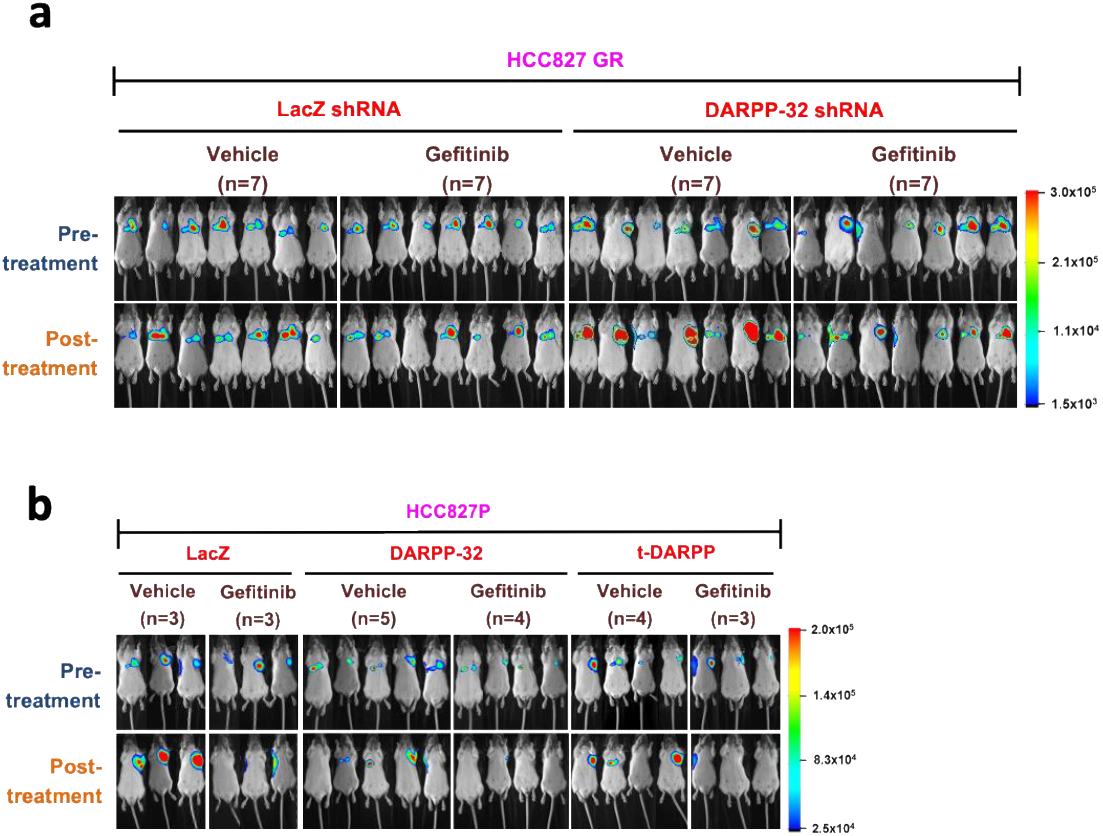
Pre- and post-treatment luminescence images of vehicle- and gefitinib-treated mice. **a** DARPP-32-depleted luciferase-labeled human HCC827GR cells were orthotopically injected into the left thoracic cavity of SCID mice. Mice administered either vehicle or gefitinib (25mg/Kg) were imaged for luminescence before and after treatment. **b** Retrovirus encoding control (LacZ), DARPP-32 or t-DARPP cDNAs were transduced in luciferase-labeled human HCC827P cells. After establishment of the tumor, mice were treated with vehicle or gefitinib (25mg/Kg) three times in a week. Luminescence images of mice were taken pre- and post-treatment. The colored bar represents the numerical value of luminescence.

**Supplementary Figure 10:**
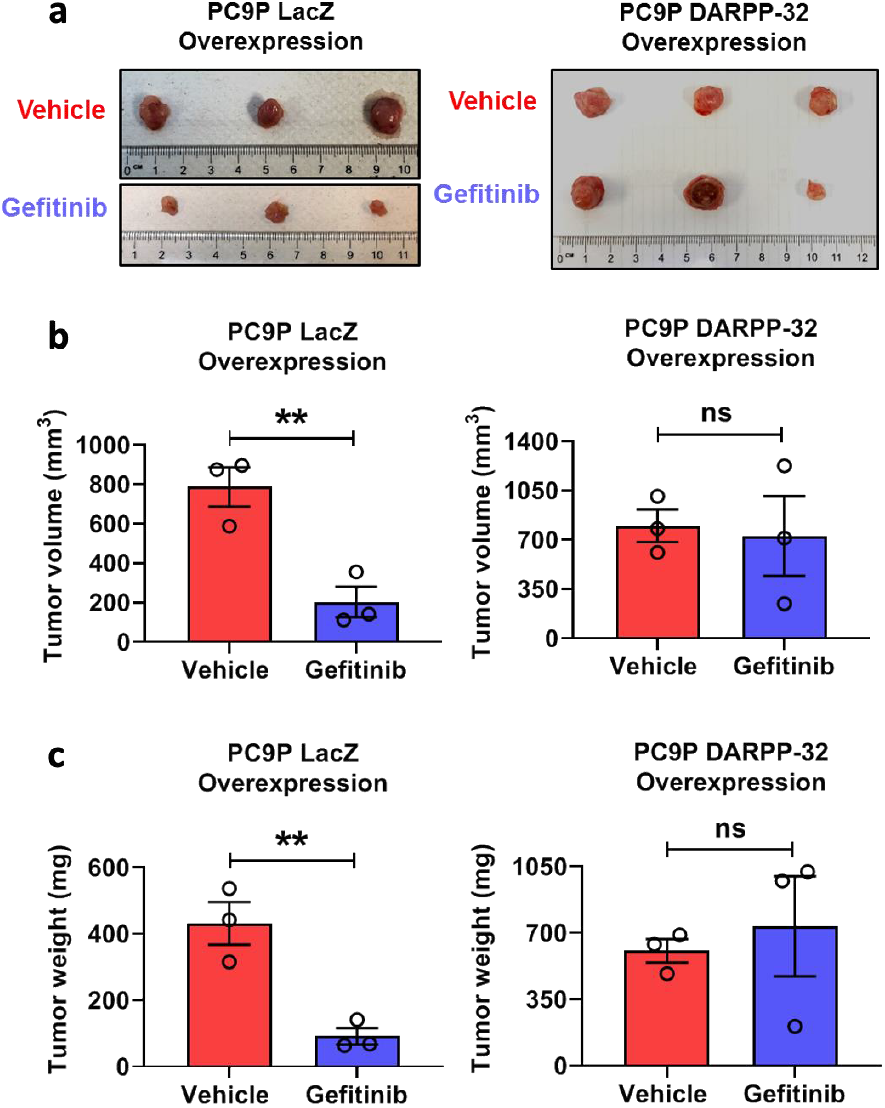
Overexpression of DARPP-32 suppresses gefitinib efficacy *in vivo*. **a-c** SCID mice were subcutaneously injected with PC9P cells stably overexpressing control (LacZ) or DARPP-32 cDNAs and treated with vehicle or Gefitinib (25mg/Kg). At endpoint of the experiments, mice were sacrificed and xenografted tumors were extirpated. Photographs of extirpated tumors were taken to visualize gross morphology (a). Tumor volume was calculated from calipers measurement after extirpation (b). Weight of extirpated tumors from sacrificed mice was measured using a digital balance (c). Each open circle on bar graphs represents an individual mouse. Bar diagrams show Mean±SEM. **P<0.01, 2-way unpaired t-test.

**Supplementary Table 1.**
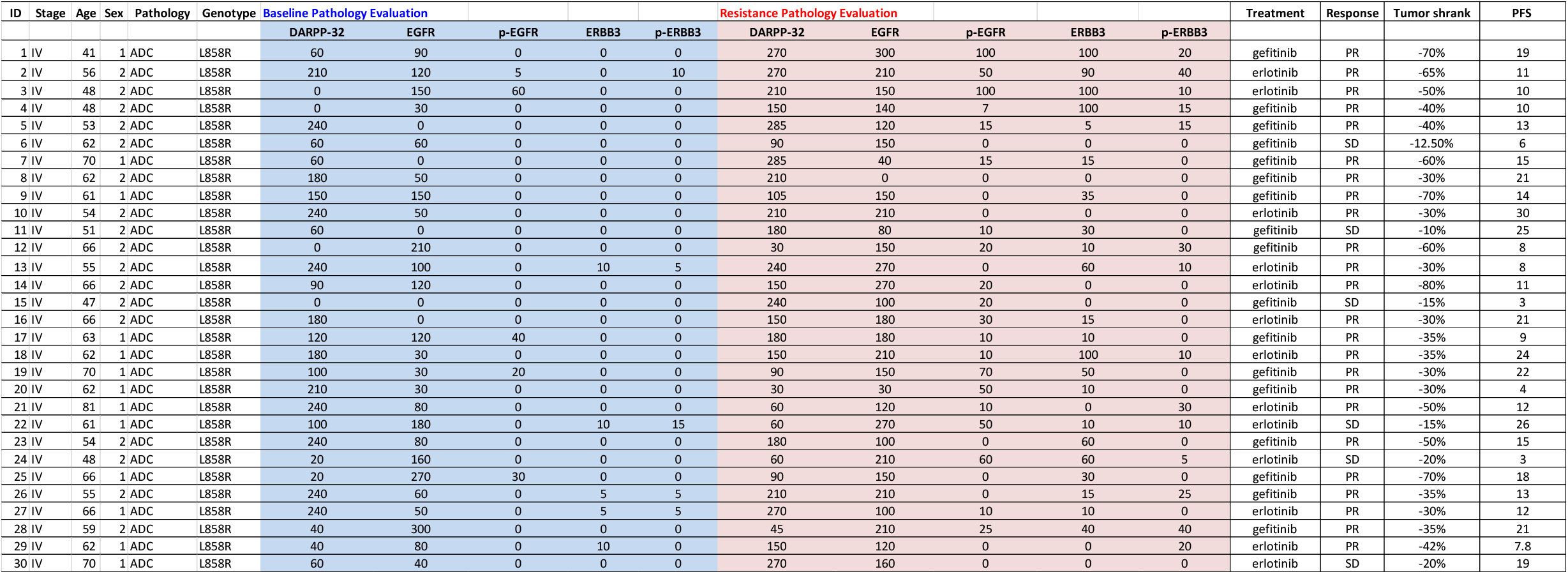

## Notes

### Competing Interest Statement

The authors have declared no competing interest.

